# Assortative mating by population of origin in a mechanistic model of admixture

**DOI:** 10.1101/743476

**Authors:** Amy Goldberg, Ananya Rastogi, Noah A Rosenberg

## Abstract

Populations whose mating pairs have levels of similarity in phenotypes or genotypes that differ systematically from the level expected under random mating are described as experiencing assortative mating. Excess similarity in mating pairs is termed positive assortative mating, and excess dissimilarity is negative assortative mating. In humans, empirical studies suggest that mating pairs from various admixed populations—whose ancestry derives from two or more source populations—possess correlated ancestry components that indicate the occurrence of positive assortative mating on the basis of ancestry. Generalizing a two-sex mechanistic admixture model, we devise a model of one form of ancestry-assortative mating that occurs through preferential mating based on source population. Under the model, we study the moments of the admixture fraction distribution for different assumptions about mating preferences, including both positive and negative assortative mating by population. We consider the special cases of assortative mating by population that involve a single admixture event and that consider a model of constant contributions to the admixed population over time. We demonstrate that whereas the mean admixture under assortative mating is equivalent to that of a corresponding randomly mating population, the variance of admixture depends on the level and direction of assortative mating. In contrast to standard settings in which positive assortment increases variation within a population, certain assortative mating scenarios allow the variance of admixture to decrease relative to a corresponding randomly mating population: with the three populations we consider, the variance-increasing effect of positive assortative mating within a population might be overwhelmed by a variance-decreasing effect emerging from mating preferences involving other pairs of populations. The effect of assortative mating is smaller on the X chromosome than the autosomes because inheritance of the X in males depends only on the mother’s ancestry, not on the mating pair. Because the variance of admixture is informative about the timing of admixture and possibly about sex-biased admixture contributions, the effects of assortative mating are important to consider in inferring features of population history from distributions of admixture values. Our model provides a framework to quantitatively study assortative mating under flexible scenarios of admixture over time.

## 1 Introduction

Mechanistic models describing the dynamics of admixture among two or more populations have proven informative for understanding the processes that underlie patterns of admixture in admixed populations. Such models have examined a variety of phenomena, including sex-biased admixture (GOLDBERG *et al*., 2014; GOLDBERG and ROSENBERG, 2015), the interaction of admixture with migration and population size (LONG, 1991; POOL and NIELSEN, 2009), the consequences of spatial structure for ancestry patterns (WANG *et al*., 2011; SEDGHIFAR *et al*., 2015), hybrid incompatibilities and epistasis (LINDTKE and BUERKLE, 2015; SCHUMER *et al*., 2015; SCHUMER and BRANDVAIN, 2016; SEDGHIFAR *et al*., 2016), and the effect of admixture on linkage disequilibrium (CHAKRABORTY and WEISS, 1988; PFAFF *et al*., 2001; GRAVEL, 2012; LOH *et al*., 2013; LIANG and NIELSEN, 2014; NI *et al*., 2016; ZAITLEN *et al*., 2017).

Recently, a general family of admixture models has introduced a framework for examining the effects of an ongoing admixture process on the properties of admixture levels in populations. VERDU and ROSENBERG (2011) devised a mechanistic model of admixture allowing for flexible admixture contributions over time, studying the dynamics of the distribution of ancestry within an admixed population. GOLDBERG *et al*. (2014) and GOLDBERG and ROSENBERG (2015) extended this model to allow for varying contributions from males and females, evaluating the consequences of sex-biased admixture for autosomal and X-chromosomal ancestry. GRAVEL (2012) derived the distribution of autosomal ancestry and ancestry-tract lengths under a related model allowing for multiple waves of migration. These studies demonstrate the potential of mechanistic admixture models for accommodating complex admixture histories.

One phenomenon of recent interest in the study of admixed populations is assortative mating, in which admixture levels of individuals in a population system that includes an admixed population and its source populations influence the formation of mating pairs. Empirical analyses of mating pairs in admixed populations have documented nonrandom pairings with respect to admixture levels (RISCH *et al*., 2009; SEBRO *et al*., 2010; ZAITLEN *et al*., 2017). In particular, ZOU *et al*. (2015) found that in multiple Latino populations, spouse pairs are correlated in their genomic ancestry patterns. Assortative mating by admixture has also been suggested as a cause of geographic structure in patterns of genetic and phenotypic variation in admixed populations (RUIZ-LINARES *et al*., 2014), and as an explanation for empirically observed differences between phenotype-based ancestry classifications and genomic ancestry (PARRA *et al*., 2003).

A particular form of assortative mating by admixture can be more specifically characterized as assortative mating by source population (RITCHIE *et al*., 1989; HOWARD, 1993; DUENEZ-GUZMAN *et al*., 2009; MELO *et al*., 2009; RUIZ-LINARES *et al*., 2014; SCHUMER *et al*., 2017). In this type of assortative mating by admixture, different groups—source populations and admixed populations, for example—come to have distinct geographic locations, trait preferences, host preferences, or in the case of human populations, social identities (QIAN, 1997; JACOBSON *et al*., 2004). Individual ancestry levels within the admixed group are less important for mating choices than the group membership itself. Positive assortative mating by source population can take place, in which individuals from the source populations preferentially mate within the source groups, while individuals from the admixed group preferentially mate within the admixed group (Figure 1b). Alternatively, negative assortative mating by source population can also take place—for example, during major migration events—in which pairings might be more likely to involve individuals from different source populations (Figure 1c).

**Figure 1:**
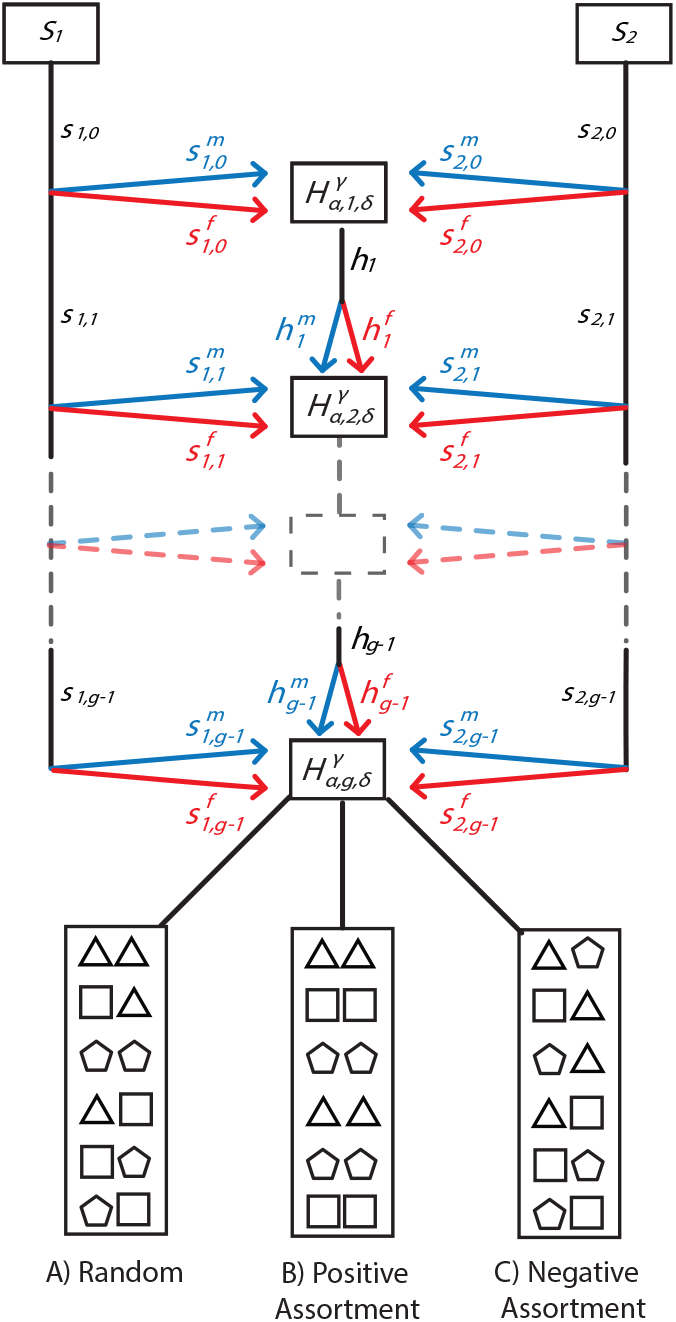
Schematic of the mechanistic model of assortative mating by population. Two source populations, ***S***_1_ and ***S***_2_, contribute females and males to the next generation of the hybrid population ***H***, potentially with time-varying proportions. The fractional contributions of the source populations and the hybrid population to the next generation ***g*** are ***s***_1,*g*_, ***s***_2,*g*_, and ***h***_*g*_, respectively. Sex-specific contributions from the populations are 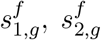 and 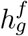 and 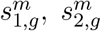 and 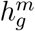, for females and males, respectively. 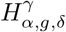 represents the fraction of admixture from source population *α* ∈ {1, 2} in generation *g* for a random individual of sex *δ* ∈ {*f*, *m*} in population *H* for chromosomal type *γ* ∈ {*A*, *X*}. Within the admixed population, at every generation, parents from generation *g* − 1 pair according to one of three mating models. Individuals from *S*_1_ are represented by triangles, *S*_2_ by pentagons, and *H* by squares. (A) Random mating. The probability of a pairing is given by the product of the proportional contributions of the two populations. (B) Positive assortative mating. Individuals are more likely to mate with individuals from their own population. (C) Negative assortative mating. Individuals are more likely to mate with individuals from a different population. In each panel, a mating pair is indicated by a pair of adjacent symbols. Each panel considers the same values for the contributions from the three populations to generation *g* + 1.

Here, we extend mechanistic models describing the evolution of admixture levels in an admixed population to accommodate assortative mating by source population. In Section 2, we develop the mechanistic admixture model, allowing mating pairs to vary in their probability of occurrence based on individual populations of origin. In Sections 3 and 4, we derive recursive expressions for the moments of autosomal and X-chromosomal admixture, respectively, as functions of sex-specific contributions and the properties of assortative mating. Next, in Sections 5 and 6, we analyze the behavior of the moments of the admixture fraction distribution for specific cases of the admixture model with constant contributions over time. We conclude with a discussion in Section 7.

## 2 Model

We follow the notation of GOLDBERG and ROSENBERG (2015), extending earlier mechanistic admixture models to consider an admixed population that preferentially mates based on population of origin. That is, individuals from each of the populations—source populations *S*_1_ and *S*_2_, and the hybrid population, *H*—preferentially mate with individuals of the other sex based on population of origin. We consider the case of two source populations, so that individuals come from one of three populations: *S*_1_, *H*, or *S*_2_.

Following GOLDBERG and ROSENBERG (2015), we define the parameters 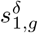 and 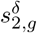 as the fractions of individuals of sex *δ* at generation *g* that trace to source populations *S*_1_ and *S*_2_, respectively, in the previous generation. For the admixed population *H*, 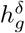 is the corresponding parameter. We maintain relations between parameter sets from GOLDBERG *et al*. (2014, eqs. 1-6). Specifically, female (superscript *f*) and male (superscript *m*) contributions must each sum to 1,

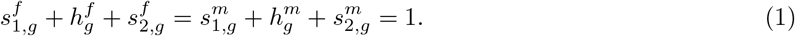

Additionally, the total contribution from a source population is the average of the sex-specific contributions from that source,

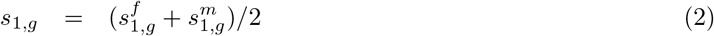

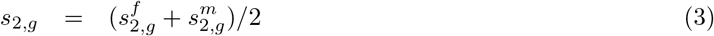

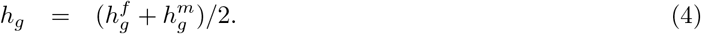

For a randomly mating population, the probability that an individual in the admixed population has a specific parental pairing is the product of the associated female and male contribution parameters. For example, the probability that an individual in the admixed population in generation *g* has a female parent from *S*_1_ and a male parent from *H* is 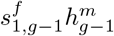. To consider deviations from random mating, we define a new parameter *c*_*ij*,*g*_ as the difference between the probability that a mating pair in generation *g* contains a female from population *i* and a male from population *j* (Table 1) and the corresponding probability in a randomly mating population with the same contribution parameters. Here, *i*, *j ϵ* {1, *h*, 2} for populations *S*_1_, *H*, and *S*_2_, respectively.

**Table 1:** Recursions for autosomal and X-chromosomal admixture. The table shows the probability of parental pairings and the admixture fractions for each of the nine possible pairings for a randomly chosen female and a randomly chosen male from the admixed population at generation *g*.

| Case | Female parent's population | Male parent's population | Probability | Admixture in males |  | Admixture in females |  |
| --- | --- | --- | --- | --- | --- | --- | --- |
| | | | | $H_{1,g,m}^A$ | $H_{1,g,m}^X$ | $H_{1,g,f}^A$ | $H_{1,g,f}^X$ |
| 1 | $S_1$ | $S_1$ | $s_{1,g-1}^f s_{1,g-1}^m + c_{11,g}$ | 1 | 1 | 1 | 1 |
| 2 | $S_1$ | $H$ | $s_{1,g-1}^f h_{g-1}^m + c_{1h,g}$ | $\frac{1 + H_{1,g-1,m}^A}{2}$ | 1 | $\frac{1 + H_{1,g-1,m}^A}{2}$ | $\frac{1 + H_{1,g-1,m}^X}{2}$ |
| 3 | $S_1$ | $S_2$ | $s_{1,g-1}^f s_{2,g-1}^m + c_{12,g}$ | $\frac{1}{2}$ | 1 | $\frac{1}{2}$ | $\frac{1}{2}$ |
| 4 | $H$ | $S_1$ | $h_{g-1}^f s_{1,g-1}^m + c_{h1,g}$ | $\frac{1 + H_{1,g-1,f}^A}{2}$ | $H_{1,g-1,f}^X$ | $\frac{1 + H_{1,g-1,f}^A}{2}$ | $\frac{1 + H_{1,g-1,f}^X}{2}$ |
| 5 | $H$ | $H$ | $h_{g-1}^f h_{g-1}^m + c_{hh,g}$ | $\frac{H_{1,g-1,f}^A + H_{1,g-1,m}^A}{2}$ | $H_{1,g-1,f}^X$ | $\frac{H_{1,g-1,f}^A + H_{1,g-1,m}^A}{2}$ | $\frac{H_{1,g-1,f}^X + H_{1,g-1,m}^X}{2}$ |
| 6 | $H$ | $S_2$ | $h_{g-1}^f s_{2,g-1}^m + c_{h2,g}$ | $\frac{H_{1,g-1,f}^A}{2}$ | $H_{1,g-1,f}^X$ | $\frac{H_{1,g-1,f}^A}{2}$ | $\frac{H_{1,g-1,f}^X}{2}$ |
| 7 | $S_2$ | $S_1$ | $s_{2,g-1}^f s_{1,g-1}^m + c_{21,g}$ | $\frac{1}{2}$ | 0 | $\frac{1}{2}$ | $\frac{1}{2}$ |
| 8 | $S_2$ | $H$ | $s_{2,g-1}^f h_{g-1}^m + c_{2h,g}$ | $\frac{H_{1,g-1,m}^A}{2}$ | 0 | $\frac{H_{1,g-1,m}^A}{2}$ | $\frac{H_{1,g-1,m}^X}{2}$ |
| 9 | $S_2$ | $S_2$ | $s_{2,g-1}^f s_{2,g-1}^m + c_{22,g}$ | 0 | 0 | 0 | 0 |

The parameters *c*_*ij*,*g*_ govern the strength and direction of assortative mating. We assume that the assortative mating preference is constant over time after the founding of the admixed population. Therefore, we have two sets of parameters: *c_ij,_*_0_ for the founding generation, and *c*_*ij*_ for all further generations. Because the sum of the probabilities for all parental pairings must be 1, we have

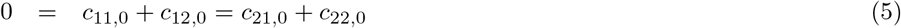

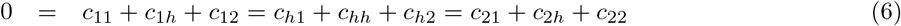

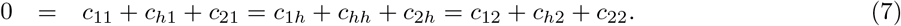

The values of the *c*_*ij*_ are bounded such that the probability of each given parental pairing (Table 1) takes its values in the interval [0, 1], and such that each probability is no greater than the probability of one of its constituent components. For example, if *c*_11_ > 0, then 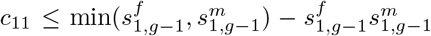. If *c*_11_ < 0, then 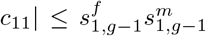. The value of *c*_*ij*_ is necessarily zero if the female contribution from population *i* or the male contribution from population *j* is zero.

For positive assortative mating in population *i*, with *i* ∈ {*S*_1_, *H*, *S*_2_}, individuals in population *i* are more likely to mate with individuals from their own population than from other populations, so that *c*_*ii*_ > 0 and *c*_*ii*_ > *c*_*ij*_ and *c*_*ii*_ > *c*_*ji*_ for each *j* ≠ *i*. If individuals from a source population are least likely to mate with individuals from the other source population, then we also have *c*_12_ *<* 0 and *c*_21_ *<* 0. Similarly, for negative assortative mating, individuals in each population are less likely to mate with individuals from their own population than with individuals from other populations, so that *c*_*ii*_ < 0 for all *i* and *c*_*ii*_ < *c*_*ij*_ and *c*_*ii*_ < *c*_*ji*_ for all *i* and *j* ≠ *i*. If individuals from a source population are most likely to mate with individuals from the other source, then *c*_12_ > 0 and *c*_21_ > 0.

We study the random variable 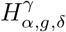 describing the admixture fraction from source population *α* at generation *g* for a chromosomal type *γ* in a random individual of sex *δ* in the admixed population. The chromosomal type is indicated by *γ*, and it can be either autosomal, *A*, or X-chromosomal, *X*. For the autosomes, 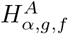 and 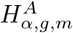 are identically distributed (GOLDBERG *et al*., 2014). For the X chromosome, the distribution of 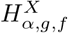 depends on both female and male admixture distributions in generation *g* − 1, as females inherit one X chromosome from each parent. For 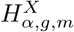, the distribution depends only on the female contributions, as males inherit a single X chromosome from their mothers.

Based on these inheritance patterns, we can write the values for the autosomal and X-chromosomal admixture fractions of an individual randomly chosen from the admixed population given one of nine possible sets of parents, *L*, along with the probability that an individual has that set of parents (Table 1). The probability of a parental pairing is a function of the sex-specific contributions from the populations and the assortative mating parameters *c*_*ij*_. If *c*_*ij*,0_ = 0 and *c*_*ij*_ = 0 for all parental pairings, with *j* not necessarily distinct from *i*, this model reduces to the random-mating model of GOLDBERG and ROSENBERG (2015).

## 3 Moments of the autosomal admixture fraction

### 3.1 Expectation

Under the model, following GOLDBERG *et al*. (2014, eq. 12), we can use the law of total expectation to write a recursion for the expected value of the admixture fraction from source population 1 for a random individual of sex *δ* sampled from the admixed population in generation *g* as a function of conditional expectations for all possible parental pairs *L*. As 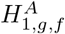 and 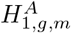 are identically distributed, we write expressions for sex *δ* when considering autosomal admixture, understanding that *δ* takes on the same value, *f* or *m*, throughout Section 3. Using the values from Table 1, for the first generation, in which neither parent is from population *H*, we have

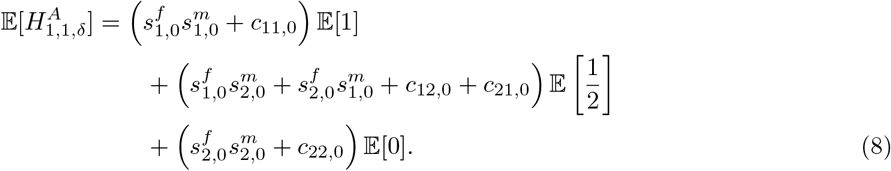

For all subsequent generations, *g* ≥ 2, we have

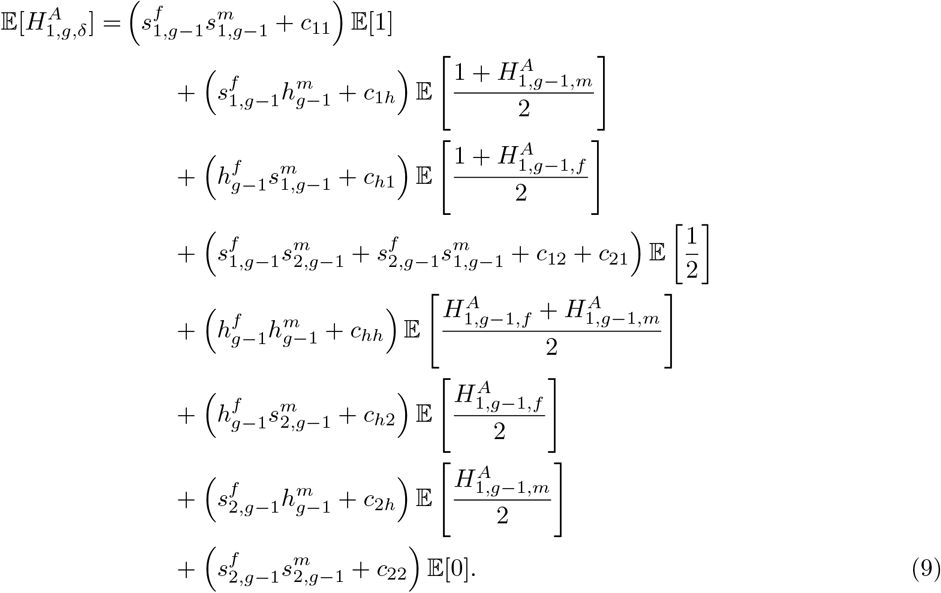

Using eqs. (1)–(4), we can simplify eq. (8) to give, for *g* = 1,

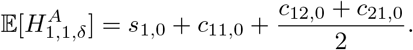

For *g* ≥ 2, recalling that 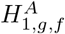 and 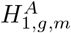 are identically distributed, eq. (9) becomes

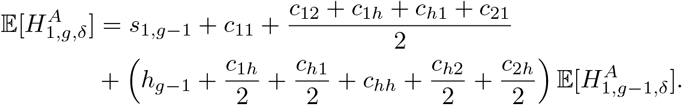

Applying eqs. (5)–(7), for the expected value of autosomal admixture in a random individual of sex *δ* from the admixed population sampled at *g* = 1, we have

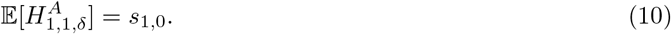

For subsequent generations *g* ≥ 2, we have

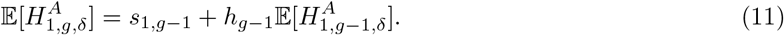

Eqs. (10) and (11) are the same as were found under a randomly mating population by GOLDBERG *et al*. (2014, eqs. 17 and 19). That is, positive or negative assortative mating according to population of origin in an admixed population does not affect the expectation of autosomal admixture. The concordance of mean ancestry under assortative and random mating makes sense because the conditions in eqs. (5)–(7) maintain the ancestry proportions from generation to generation in a manner that does not permit natural selection favoring one or another ancestral membership. The ancestries of the parents contributing to the next generation are the same in both scenarios; assortative mating only changes the probability distribution of the parental pairing.

### 3.2 Higher moments

Next, we write a general recursion for the higher moments of the autosomal admixture fraction from population *S*_1_ in a randomly chosen individual of sex *δ* from the admixed population. For moments *k* ≥ 1, similar to eqs. 20 and 21 of GOLDBERG *et al*. (2014) in generation *g* = 1, we obtain

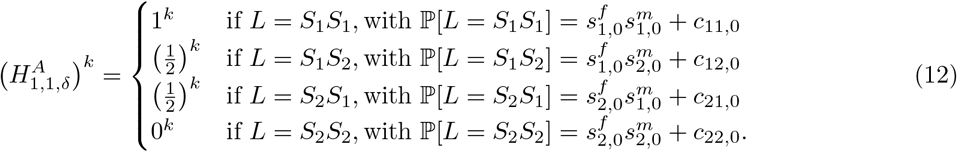

For *g* ≥ 2, we have

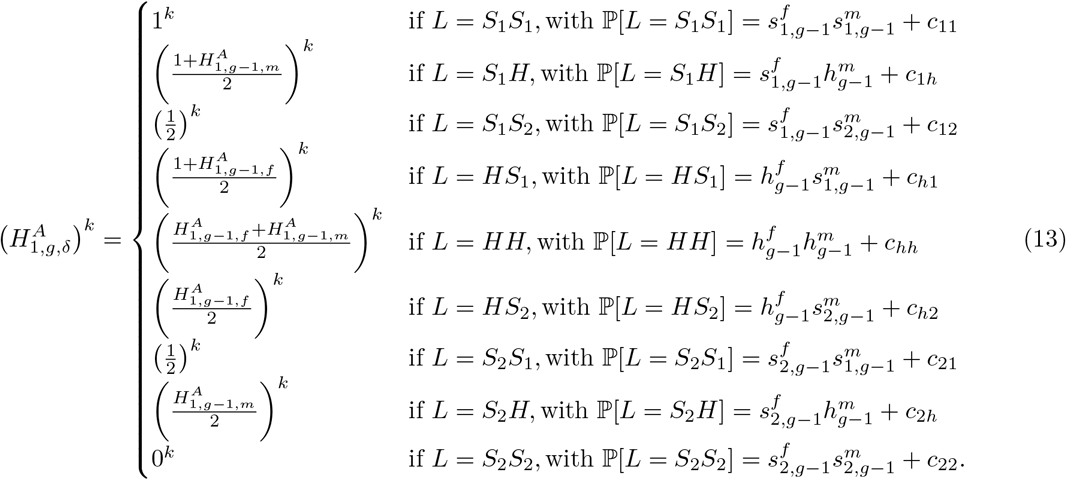

For *k* = 1, eqs. (12) and (13) simplify to produce the recursion in Table 1. Using the law of total expectation, and following our calculation for the expected value of autosomal admixture, we can write an expression for the higher moments of autosomal admixture by summing the conditional values for autosomal admixture given the parental pairings over all possible sets of parental source populations. For the first generation, *g* = 1, we have

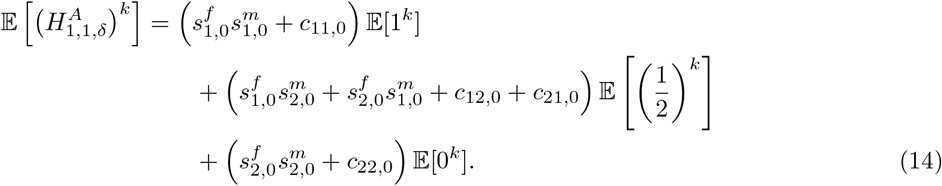

For *g* ≥ 2, we have

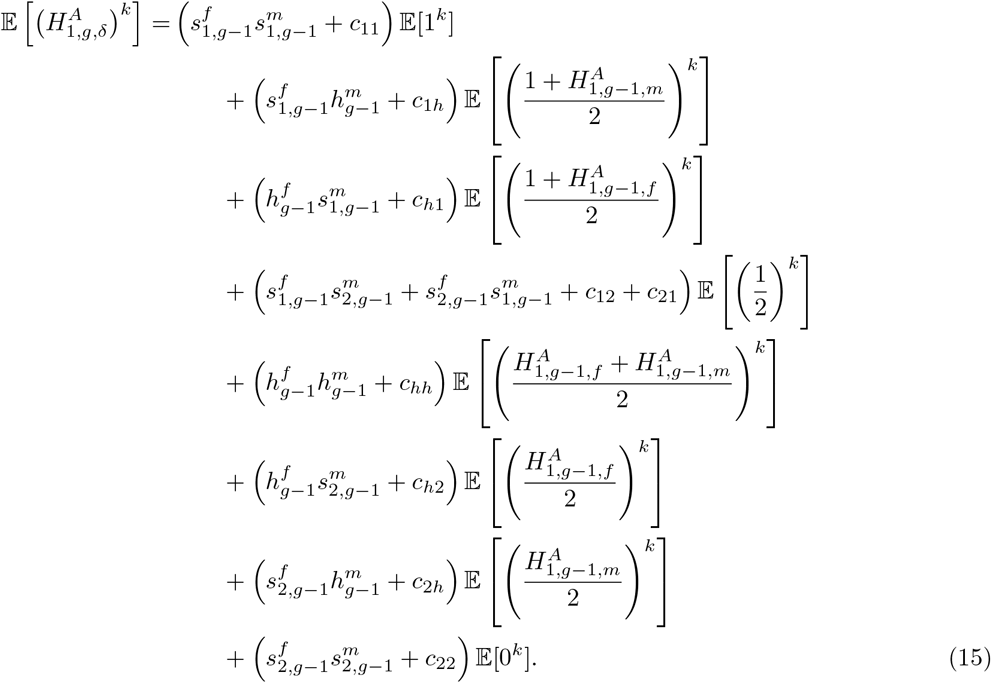

Recalling that 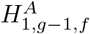 and 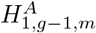 are conditionally independent given 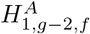 and 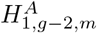 (GOLDBERG *et al*., 2014), we use the binomial theorem to simplify eqs. (14) and (15). For *g* = 1, we have

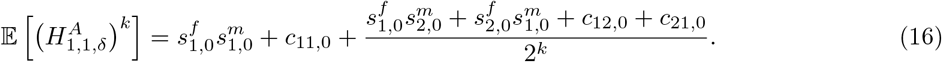

For *g* ≥ 2, we have

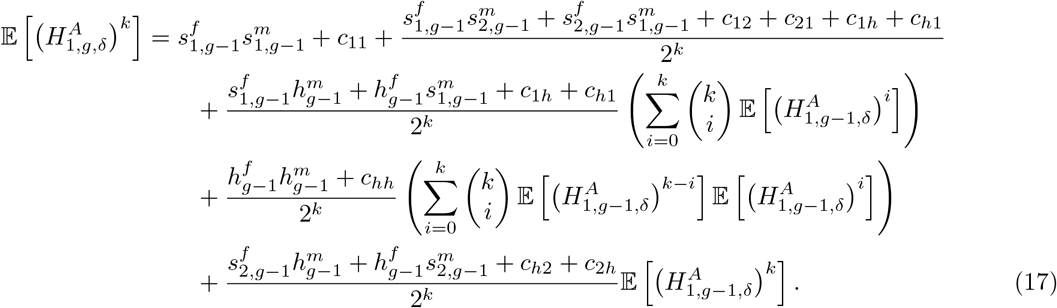

For *k* = 1, eqs. (16) and (17) reduce to eqs. (10) and (11).

The recursions for the higher moments of the autosomal admixture fraction distribution follow the corresponding expressions for a randomly mating population, but with additional terms that are linear in the coefficients for the mating preferences. Setting the assortative mating parameters *c*_*ij*,0_ and *c*_*ij*_ equal to 0 for all *i* and *j*, eqs. (16) and (17) reduce to the recursion for the moments of admixture for a corresponding randomly mating population, eqs. 24 and 26 from GOLDBERG *et al*. (2014).

### 3.3 Variance

Using eqs. (16) and (17) for *k* = 2, we can write expressions for the second moment of autosomal admixture. Recalling eqs. (1)–(4), for *g* = 1, we have

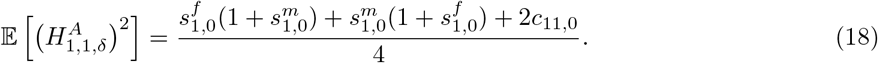

For *g* ≥ 2, because 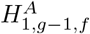 and 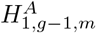 are identically distributed, we have

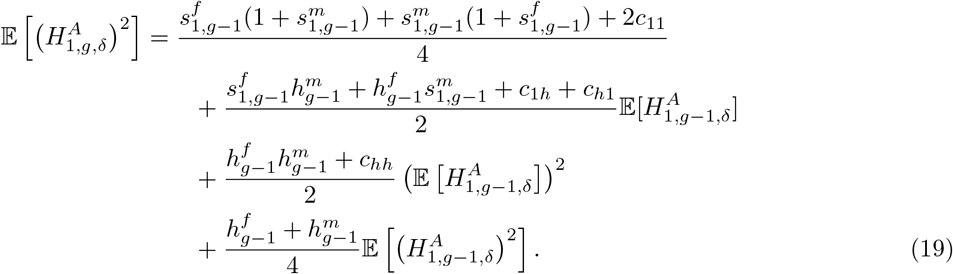

Using the definition of the variance 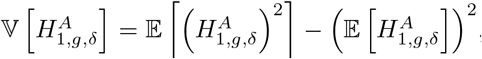, with the expressions for the expected value of autosomal admixture, eqs. (10) and (11), and the second moment, eqs. (18) and (19), we can write an expression for the variance of autosomal admixture. For *g* = 1, we have

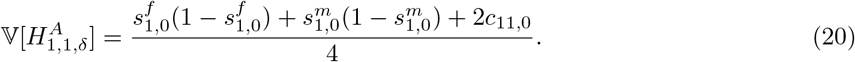

For all subsequent generations, *g* ≥ 2, we have

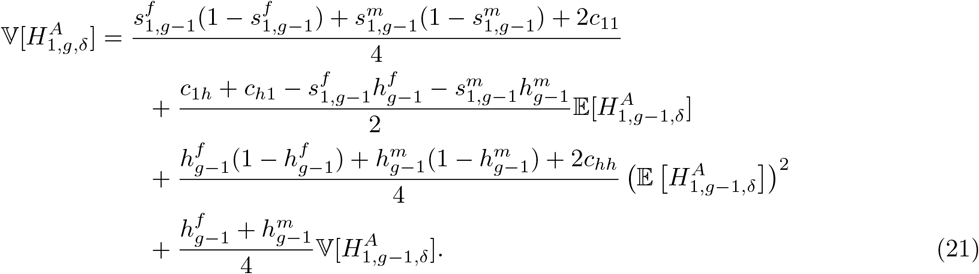

For a randomly mating population, with *c*_*ij*,0_ = 0 and *c*_*ij*,*g*_ = 0 for all *i* and *j*, eqs. (20) and (21) reduce to eqs. 32 and 33 from GOLDBERG *et al*. (2014). Defining the variance of autosomal admixture in a randomly mating population at *g* = 1 and *g* ≥ 2, eqs. 32 and 33 from GOLDBERG *et al*. (2014), as 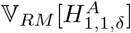 and 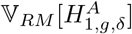, respectively, we can rewrite eqs. (20) and (21). For *g* = 1, we have

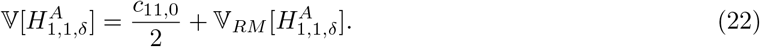

For *g* ≥ 2, recalling eqs. (6) and (7), we have

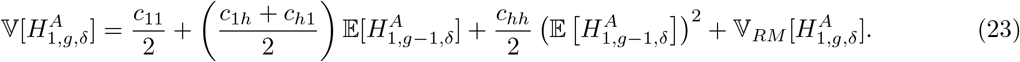

Eqs. (22) and (23) indicate that, in a single generation, assortative mating changes the variance of admixture compared to a randomly mating population. The variance of autosomal admixture contains information about assortative mating because it depends on the assortative mating parameters *c*_*ij*_ and it differs from the expression for a randomly mating population with the same contributions for the source populations.

## 4 Moments of the X-chromosomal admixture fraction

### 4.1 Expectation

Following the same approach as in the corresponding derivation for the autosomal admixture fraction, we can use the law of total expectation to write recursions for the moments of X-chromosomal admixture. Because the distribution of admixture differs for females and males on the X chromosome (Table 1), we follow GOLDBERG and ROSENBERG (2015) and write coupled expressions for the moments of female and male X-chromosomal admixture fractions.

For X-chromosomal admixture in a randomly chosen female from the admixed population, the recursive expressions for the moments of admixture match the expressions for autosomal admixture presented in Section 3, exchanging *γ* = *A* for *γ* = *X* in the superscript. However, X-chromosomal admixture is not identically distributed in females and in males from the admixed population. Therefore, certain expressions for the moments of admixture cannot be reduced as they were for the autosomes.

Using eqs. (1)–(7), we can use the values from Table 1 to write expressions for the expectation of X-chromosomal admixture. For X-chromosomal admixture in a female from the admixed population, the recursive expressions for the expectation of admixture are the same as for autosomal admixture (eqs. (10) and (11)), exchanging *γ* = *A* for *γ* = *X* in the superscript. For X-chromosomal admixture in males, for *g* = 1, we have

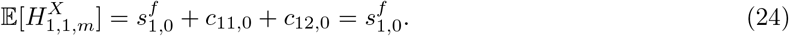

For *g* ≥ 2, we have

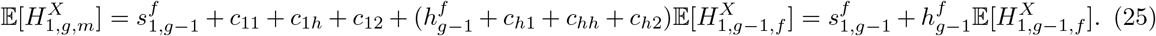

As was the case for the autosomes, the expectation of X-chromosomal admixture in eqs. (24) and (25) is the same under assortative mating as the expected value for a randomly mating population (GOLDBERG and ROSENBERG, 2015, eqs. 3–4).

### 4.2 Higher moments

Following the derivation for the autosomes and using Table 1, we can write general coupled recursions for the higher moments of the X-chromosomal admixture fraction from population *S*_1_ in a randomly chosen female and male from the admixed population. As was true for the expectation, the recursion for the X-chromosomal admixture fraction in a female from the admixed population for *k* ≥ 1 is the same as the recursion for autosomal admixture in eqs. (12) and (13), exchanging the superscript *A* for *X*. For X-chromosomal admixture in a male, we have, for *g* = 1,

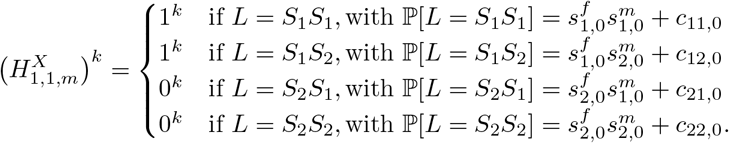

For *g* ≥ 2, we have

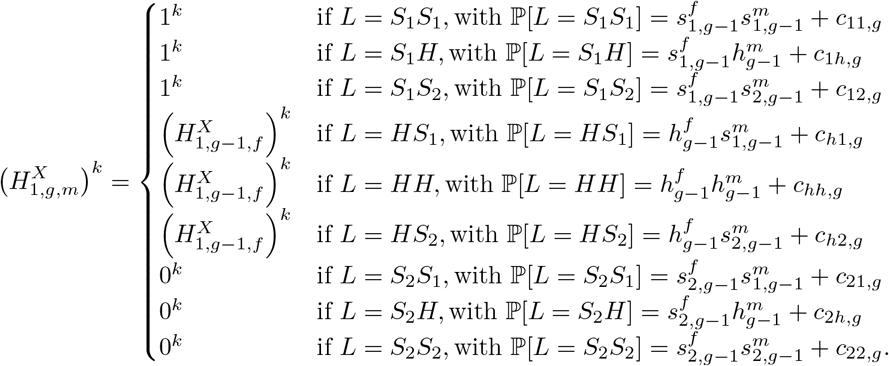

Using the law of total expectation and the binomial theorem, and following the derivation of the autosomal admixture moments, we can write simplified coupled expressions for the moments of X-chromosomal admixture, separately considering a female and a male from the admixed population for *k* ≥ 1. For *g* = 1, the recursion for the *k*th moment of X-chromosomal admixture in a female is equal to the corresponding recursion in the case of autosomal admixture, eq. (16). For a male in *g* = 1, we have

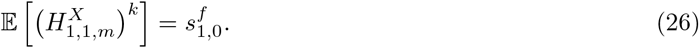

For *g* ≥ 2, using the conditional independence of 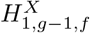 and 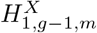 given 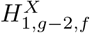 and 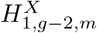, we have

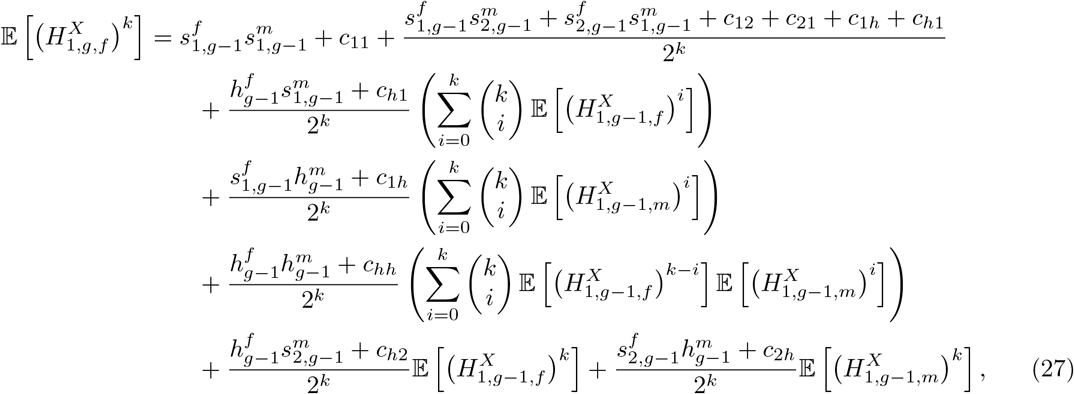

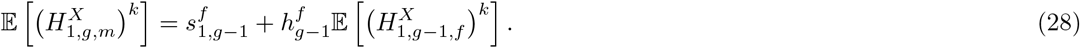

Unlike for the autosomes, the female and male admixture fractions are not identically distributed or conditionally independent, so the dependence on both 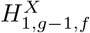 and 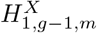 in eq. (27) cannot be further reduced. For *k* ≥ 2, moments of the X-chromosomal admixture fraction depend on the assortative mating parameters, *c*_*ij*,0_ and *c*_*ij*_. However, conditional on 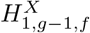, moments of the X-chromosomal admixture fraction sampled in a male from the admixed population do not depend on the assortative mating parameters. Because a single copy of the X chromosome is inherited from mother to son, the distribution of admixture for male X chromosomes in a given generation is affected only by the origin of the mother and not by the probabilities of parental pairings in Table 1.

### 4.3 Variance

For *k* = 2, we can use eqs. (26)–(28) to write coupled expressions for the second moment of X-chromosomal admixture in a randomly selected female and male from the admixed population. Recalling eqs. (1)–(4), for *g* = 1, the second moment of X-chromosomal admixture in a female is the same as the expression for autosomal admixture in eq. (18), substituting superscript *X* in place of *A*. For the second moment of X-chromosomal admixture in males in *g* = 1, we have

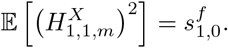

For *g* ≥ 2, we have

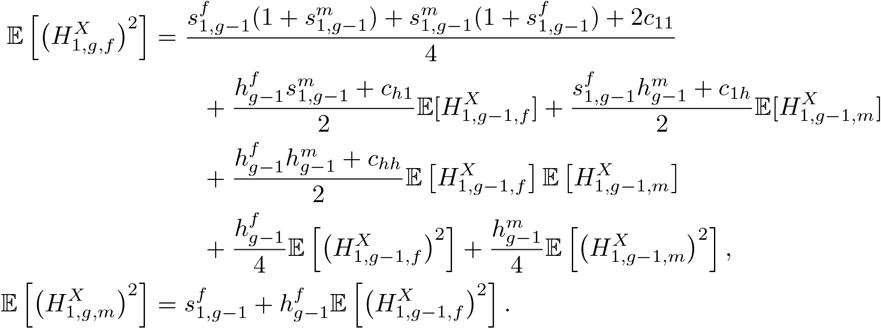

Following our derivation of the variance of autosomal admixture in Section 3.3, we can write the variance of X-chromosomal admixture in a female and in a male from the admixed population using the expected values and second moments of X-chromosomal admixture. For *g* = 1, we have

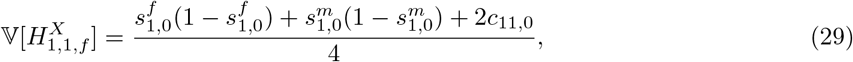

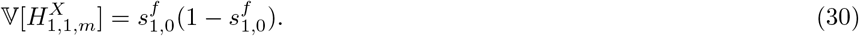

For *g* ≥ 2, we have

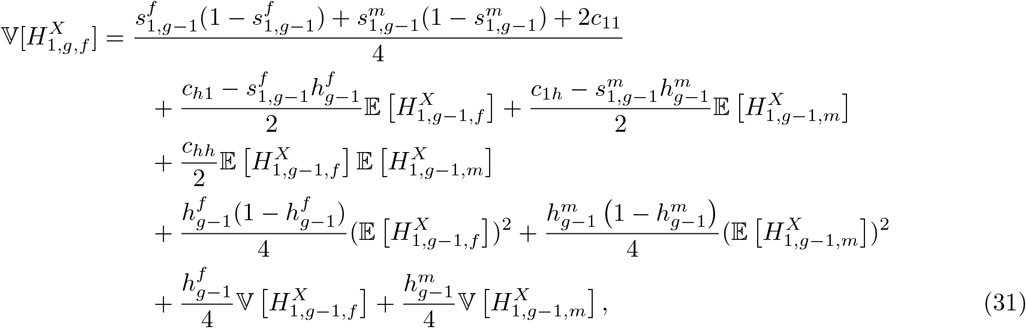

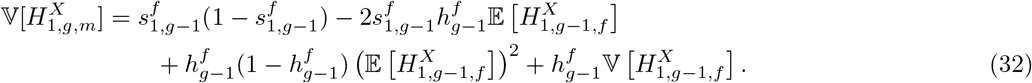

We denote the variance of X-chromosomal admixture under random mating, in a random female and male from the admixed population, by 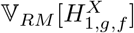 and 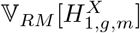 respectively. Setting all *c*_*ij*,0_ and *c*_*ij*_ parameters to zero, for *g* = 1, we have

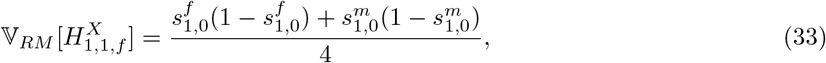

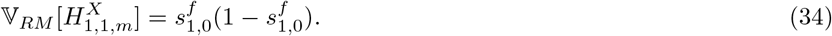

For *g* ≥ 2, we have

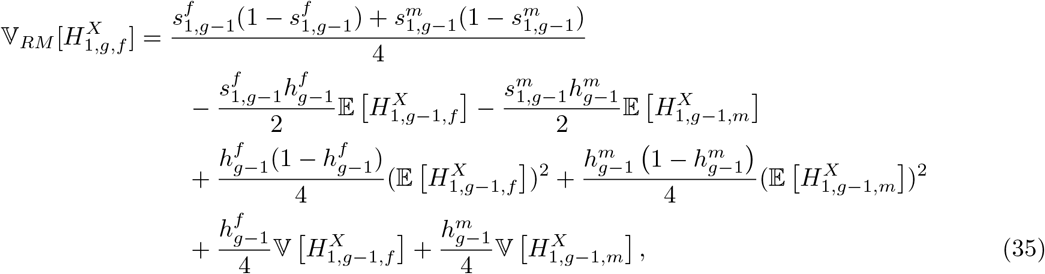

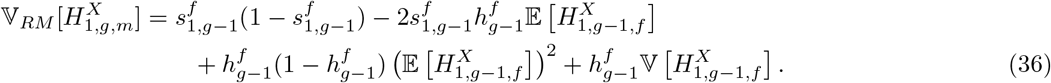

Using eqs. (33)–(36), we can rewrite the variances in eqs. (29)–(32) as functions of the variance under a similar randomly mating population. For *g* = 1, we have

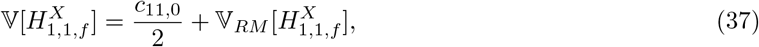

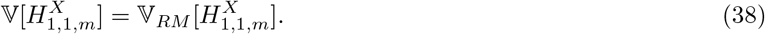

For *g* ≥ 2, conditional on 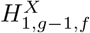 and 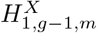, and recalling eqs. (6) and (7), we have

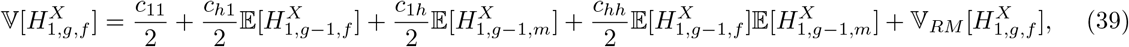

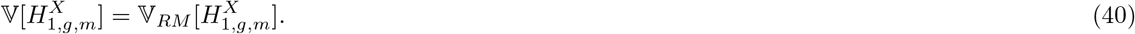

In contrast to the expectation, the variance of X-chromosomal admixture depends on the assortative mating parameters *c*_11_, *c*_*h*1_, *c*_1*h*_, and *c*_*hh*_. Conditional on the female and male variances of X-chromosomal admixture in generation *g* − 1, the variance of X-chromosomal admixture in a male sampled in generation *g* is equivalent to that in a corresponding randomly mating population.

## 5 Special case: a single admixture event

To analyze the behavior of the model, we study the moments of the distribution of autosomal and X-chromosomal admixture for two special cases of constant admixture processes over time. First, in Section 5, we consider a case with no contributions from the sources after the initial founding. That is, for *g* ≥ 2, we set 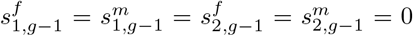 and 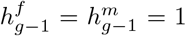. Because only the admixed population contributes after the first generation, all assortative mating parameters for further generations equal 0, *c*_*ij*_ = 0.

Next, in Section 6, we examine the special case of constant non-zero contributions over time. In Sections 5 and 6, we assume that the assortative mating process is the same for both sexes. That is, *c*_*ij*,0_ = *c*_*ji*,0_ and *c*_*ij*_ = *c*_*ji*_. The behavior of the expectation of autosomal and X-chromosomal admixture is equivalent to that seen in a randomly mating population, as studied by GOLDBERG *et al.* (2014) and GOLDBERG and ROSENBERG (2015). Therefore, as in GOLDBERG *et al.* (2014), we focus on the variance, on which the assortative mating parameters have an impact. In Section 5, we consider the variances of autosomal and X-chromosomal admixture as functions of 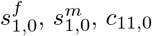, and *g*. We focus on the dependence of the variance of admixture on the assortative mating parameter *c*_11,0_. Note that the variance of admixture under random mating (*c*_11,0_ = 0) in a single-admixture scenario was studied in detail for autosomes by GOLDBERG *et al.* (2014); for completeness, we include analogous calculations for the X chromosome in Appendix A.

### 5.1 Autosomes

Under this model of a single admixture event, we can write an exact expression for the variance of autosomal admixture. Using the equation for the variance of autosomal admixture in eq. (20), we observe that the variance for a single admixture event can be written as a geometric sequence with ratio 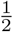. For *g* ≥ 1, we have

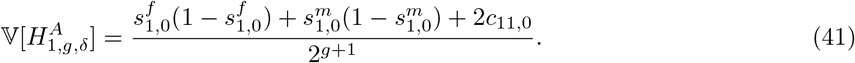

Recalling the variance of autosomal admixture for a randomly mating population produced by a single admixture event from GOLDBERG *et al.* (2014, eq. 35), we rewrite eq. (41) as a function of 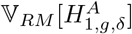,

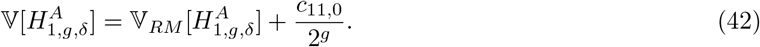

As is true in a randomly-mating population, the long-term limit of the variance over time is zero, with the effect of assortative mating being multiplied by 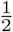 each generation, and with the distribution of admixture narrowing around the same mean value as in a randomly mating population, *s*_1,0_ (GOLDBERG *et al.*, 2014, eq. 35).

For positive assortative mating, with *c*_11,0_ > 0, the variance (eq. (42)) is larger than in a corresponding randomly mating population (GOLDBERG *et al.*, 2014, eq. 35). For negative assortative mating, *c*_11,0_ < 0, the variance is smaller (Figure 2a). The effect of the initial non-random mating on the population decreases monotonically each generation, as all individuals mate randomly within the admixed population, with no further contributions from the sources. In a given generation and given the contributions 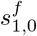 and 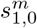, the variance is directly related to *c*_11,0_. Therefore, the maximal and minimal variance occur when *c*_11,0_ is maximized and minimized, respectively.

**Figure 2:**
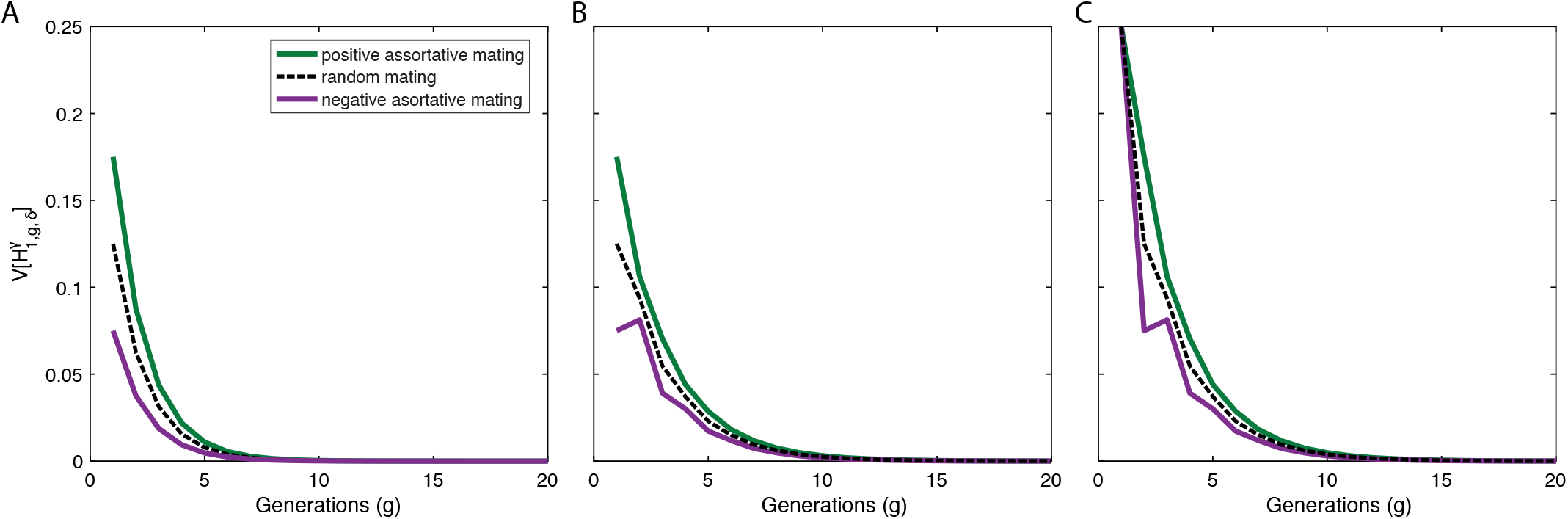
Variance of ancestry under a single-admixture scenario with assortative mating in the founding generation. (A) Autosomes,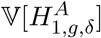. (B) X chromosomes in a female, 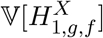. (C) X chromosomes in a male, 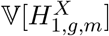. Each panel considers positive assortative mating (green), *c*_11,0_ = 0.1, negative assortative mating (purple), *c*_11,0_ = −0.1, and random mating (black), *c*_11,0_ = 0. The plots use eqs. (41), (43), and (44) with 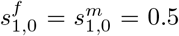. For positive assortment, the variance is higher than for random mating. For negative assortative mating, the variance is lower.

Negative assortative mating also introduces a minimum in the variance of autosomal admixture, with 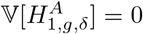, when the numerator of eq. (41) is zero, at 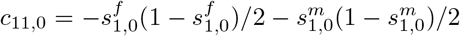 Under this scenario of complete negative assortative mating, individuals mate only with individuals from the other source population.

### 5.2 X chromosome

Similarly to the case of autosomal admixture, we can write an exact solution for the variance of X-chromosomal admixture for a scenario with a single admixture event and no subsequent admixture. We follow the derivation of the expectation of X-chromosomal admixture in GOLDBERG and ROSENBERG (2015, eqs. 6–11). Recalling eqs. (29) and (30), we rewrite the variance of X-chromosomal admixture under a scenario of a single admixture event as a coupled pair of recursions for 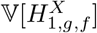 and 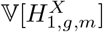.

For *g* = 1,

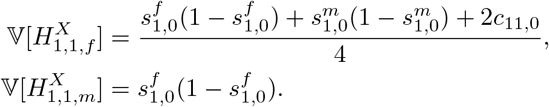

For *g* ≥ 2,

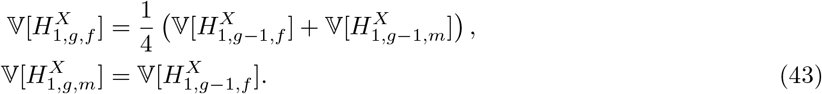

Next, for *g* ≥ 3, we can rewrite the variance of X-chromosomal admixture in a female as a two-generation recursion,

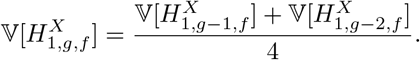

The variance of X-chromosomal admixture in a female is similar in form to the expectation seen in GOLDBERG and ROSENBERG (2015, eq. 8), and we can take an analogous approach to solving the recursion. The recursion for the variance includes a factor of 4 in each generation; therefore, the closed-form expression contains a factor of 4^*g*^. We define 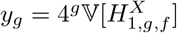 and for *g* ≥ 3, we have

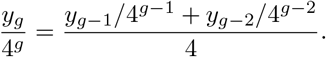

Multiplying both sides by 4^*g*^, we have a recursion 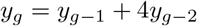, with

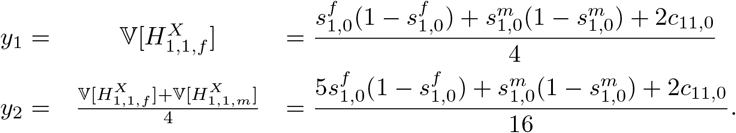

Next, for *g* ≥ 3, we write *y*_*g*_ as 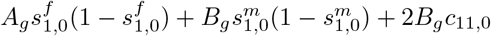, where

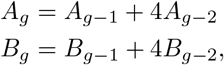

and *A*_1_ = 1, *A*_2_ = 5, *B*_1_ = 1, and *B*_2_ = 1. Calculating further values of *B*_*g*_, we note that *B*_*g*_ = *A*_*g*−1_. We can therefore rewrite 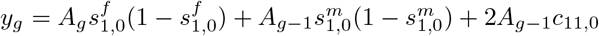, leading to

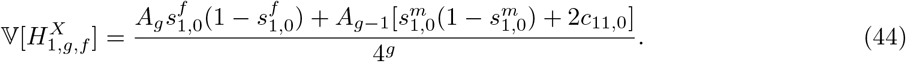

The sequence *A*_*g*_ is reminiscent of a similar recursively defined sequence that appears in the X-chromosomal expectation for a single admixture event in GOLDBERG and ROSENBERG (2015). *A*_*g*_ is a Lucas sequence (OEIS A006131), which can be written in closed form by using its generating function, *a*(*x*) = 1/(1 − *x* − 4*x*^2^). For *g* ≥ 1, we have

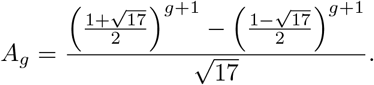

For a single admixture event, the variance of the admixture fraction in both males and females (eqs. (43) and (44)) decays to zero over time, as 4^*g*^ grows faster than *A*_*g*_. The effect of assortative mating on the X-chromosomal variance decreases at a slower rate—*c*_11,0_ accumulates a factor of 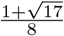 each generation—than for the autosomal variance, in which *c*_11,0_ is multiplied by a factor of 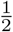 each generation (eq. (42)).

For specified contribution parameters 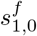 and 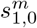 given *g*, the maximum and minimum of the variance of X-chromosomal admixture as functions of *c*_11,0_ occur at the same values of *c*_11,0_ as the maximum and minimum of the autosomal variance: the maximum and minimum of *c*_11,0_.

Figures 2–4 analyze the behavior of the variances of autosomal and X-chromosomal admixture for the case of a single admixture event. Figure 2 plots 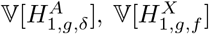 and 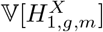 in relation to the time since admixture. For positive assortative mating, we observe that for both the autosomes (Fig. 2a) and the X chromosome (Figs. 2b and 2c), the variance of admixture is greater relative to corresponding randomly mating populations with the same contribution parameters. For negative assortative mating, the variance is smaller.

Figure 3 notes a special case of the minimum of the autosomal variance at constant 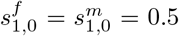 and 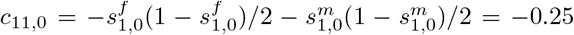. Using eq. (41), 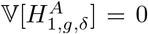 even though this set of contributions 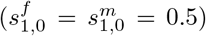 is a maximum of the variance for a randomly mating population (GOLDBERG *et al*., 2014).

**Figure 3:**
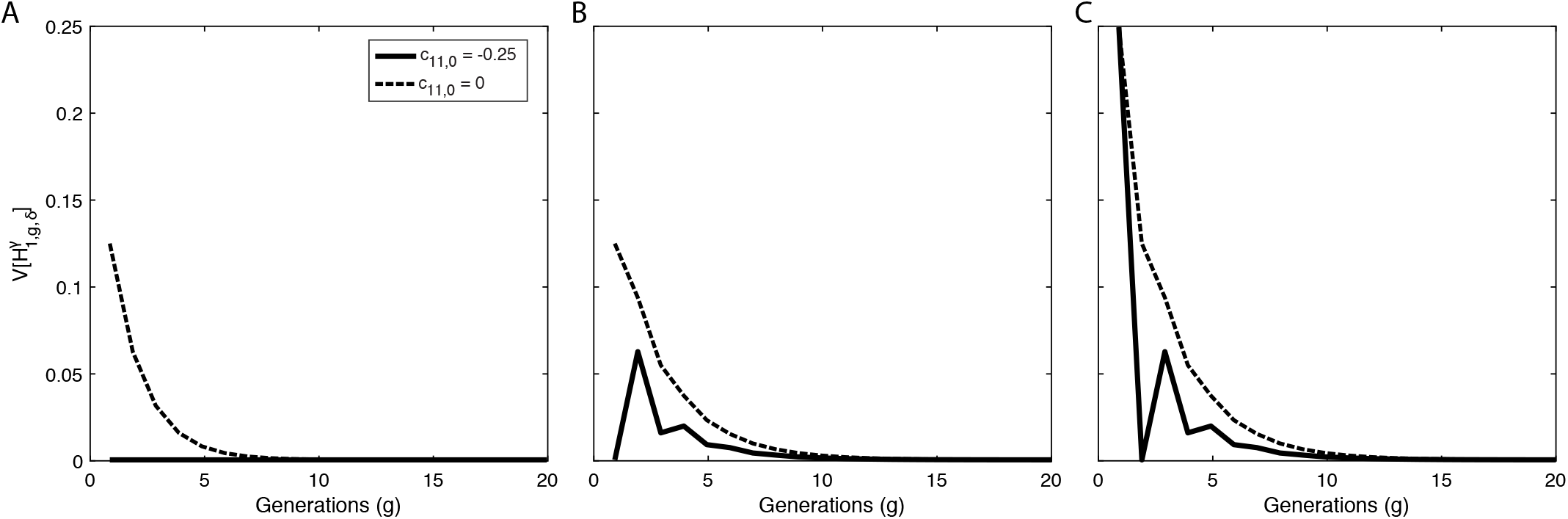
Special case of the variance of ancestry under a single-admixture scenario with negative assortative mating in the founding generation. (A) Autosomes, 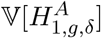. (B) X chromosomes in a female, 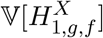. (C) X chromosomes in a male, 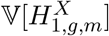. In each panel, the variance is plotted for negative assortative mating, *c*_11,0_ = −0.25, using eqs. (41), (43), and (44) with 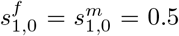. The plot highlights a special case in which the variance of autosomal admixture is zero because the numerator of eq. (41) is zero, even though 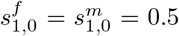 is a maximum of the variance with respect to the sex-specific contributions in a randomly mating population. The *c*_11,0_ = 0 case is copied from Figure 2.

Figure 4 plots the variance of admixture over the permissible range of *c*_11,0_, [−0.25, 0.25], for each value of *g* in [1, 6] when 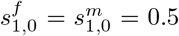. The variance of admixture increases linearly with *c*_11,0_. That is, positive values of *c*_11,0_ produce a higher variance of admixture than that of a randomly mating population (*c*_11,0_ = 0), and negative values of *c*_11,0_ produce a lower variance. The autosomal variance decreases monotonically with *g* for each value of *c*_11,0_. The X-chromosomal variance is not monotonic as *g* increases for a specific value of *c*_11,0_, as can be seen from the existence of points of intersection of the lines (Fig. 4b), and as is described for the expectation when *c*_11,0_ = 0 by GOLDBERG and ROSENBERG (2015). For the autosomes, the minimum observed in Figure 4a across values of *c*_11,0_ with 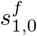 and 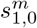 held constant occurs at *c*_11,0_ = −0.25, the minimum depicted in Figure 3, with 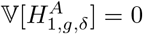 for all *g*. The male X-chromosomal admixture in *g* = 1 is 0 and does not depend on *c*_11,0_ (Fig. 4c).

**Figure 4:**
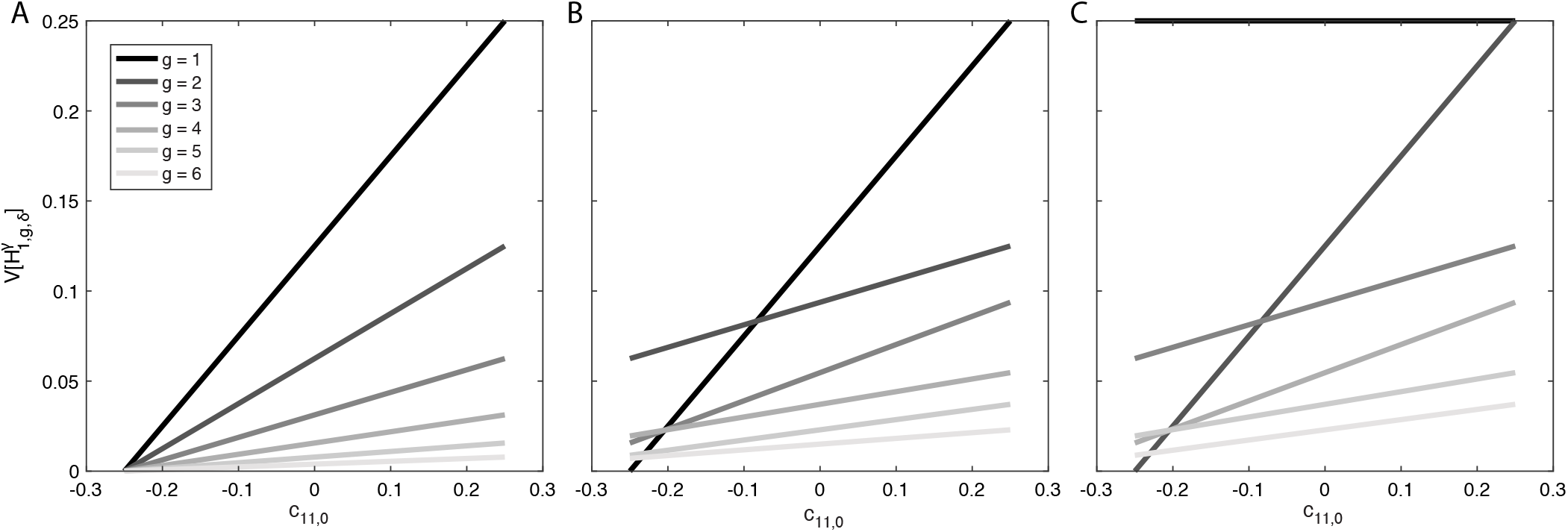
Variance of ancestry under a single-admixture scenario as a function of the assortative mating parameter *c*_11,0_. (A) Autosomes, 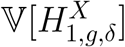. (B) X chromosomes in a female, 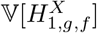. (C) X chromosomes in a male, 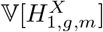. In each panel, the variance is plotted over the range of *c*_11,0_, [−0.25, 0.25], for values of *g* between 1 and 6 (from darkest to lightest gray), with 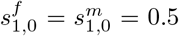, using eqs. (41), (43), and (44).

For all initial contributions 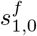 and 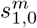, and assortative mating parameter values *c*_11,0_, the limit of the autosomal and X-chromosomal variances as *g*→∞ equals zero. That is, for admixture processes with no contributions after the founding, in both assortative and randomly mating populations, the distribution of admixture narrows around the mean.

## 6 Special case: constant contributions over time

Next, we consider the special case of constant, non-zero contributions from the source populations to the admixed population over time, after its initial founding. As in GOLDBERG *et al*. (2014) and GOLDBERG and ROSENBERG (2015), we choose the sex-specific parameters to have values that are constant in time, 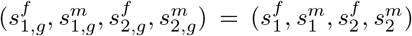 for all g ≥ 1. Sex-specific contributions take their range in [0, 1]; however, the total contributions from each source, *s*_1_ and *s*_2_, take their range in (0, 1). We maintain separate parameters for the founding contributions, 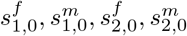. Here, we analyze the limiting behavior of the variance of autosomal and X-chromosomal admixture in an assortatively mating population under this scenario of constant contributions from the source populations to the admixed population in each generation after founding.

### 6.1 Limit of the variance of admixture over time

Appendix B uses the closed-form expressions for the variance of autosomal admixture in a randomly mating population, derived by GOLDBERG *et al*. (2014), to write a closed-form expression for the variance of autosomal admixture in eqs. (20) and (21). Taking the limit as *g* → *∞* of the expression in eq. (57) and assuming *c*_*ij*_ = *c*_*ji*_, we have

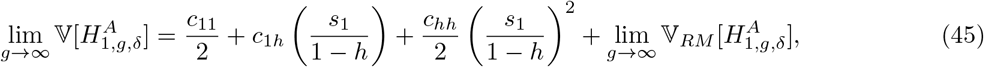

where 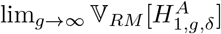 is given by GOLDBERG *et al*. (2014, eq. 53) and is reproduced in Appendix B, eq. (58).

Next, we consider the variance of X-chromosomal admixture. Using a generating function approach, Appendix C calculates the limits of 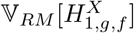 and of 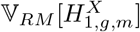 (eqs. (71) and (72)) as *g → ∞*. Recalling eqs. (37)–(40) and using eq. (73) for the definition of *P*_3_, we have

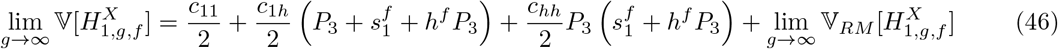

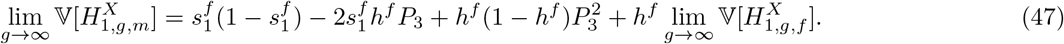

Figure 5 plots the variance of autosomal and X-chromosomal admixture over time for various values of the assortative mating parameters, both positive and negative, for constant contributions from the source populations over time. In these plots of the variance of admixture over time, we hold *c*_11_ = *c*_22_ and *c*_1*h*_ = 0, and by consequence, *c*_*h1*_ = *c*_*h2*_ = *c*_*2h*_ = 0. This set of values for the *c*_*ij*_ can be interpreted as representing (1) equal levels of within-population preference for the two source populations, and (2) for each source, the mating probability between hybrid and source individuals is the same as in a randomly mating population.

**Figure 5:**
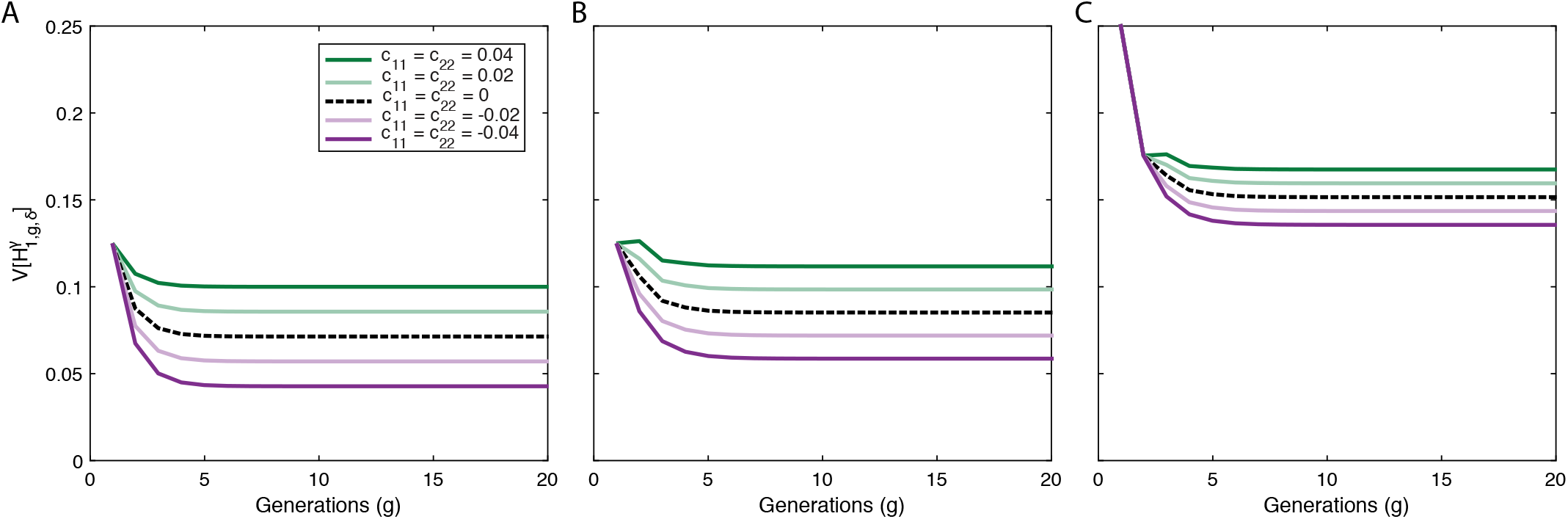
Variance of ancestry under a constant-admixture scenario with assortative mating in producing *g* ≥ 2. (A) Autosomes, 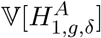. (B) X chromosomes in a female, 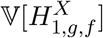. (C) X chromosomes in a male, 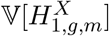. Each panel considers positive assortative mating (greens), negative assortative mating (purples), and random mating (black dashed). The plots use eqs. (20), (21), and (29)–(32). We fix *c*_11_ =*c*_22_ and *c*_1*h*_ = 0, with 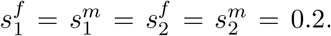. For initial conditions, 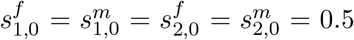 and *c*_11,0_ = 0. Positive assortative mating increases the variance relative to a randomly mating population, whereas negative assortative mating decreases it.

These limiting variances often differ from those of corresponding randomly mating populations with the same contributions. For the autosomes (Fig. 5a) and the X chromosome (Figs. 5b,c), the variance has a linear relationship with *c*_*ii*_, with positive values increasing the variance and negative values decreasing it. The limits of the autosomal and X-chromosomal variances in admixture depend on the ongoing sex-specific contributions from the source populations, 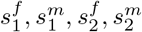 and on the assortative mating parameters *c*_*ij*_. The limits do not depend on the contributions or assortative mating parameters in the founding generation, 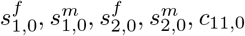.

### 6.2 Positive assortative mating can increase or decrease the variance relative to a corresponding randomly mating population

We next consider the effect of assortative mating on the variance of admixture compared to a corresponding randomly mating population with the same contribution parameters. We demonstrate that the behavior of the variance depends on the nature of the assortative mating parameters. That is, positive assortative mating can either increase or decrease the variance relative to a corresponding randomly mating population.

First, we consider assortative mating in which preference for one’s own population is the largest of the three preferences, and, for source populations, preference for the other source population is the smallest of the three. Additionally, all three populations, *S*_1_, *H*, *S*_2_, experience the same direction of mating preference. For positive assortative mating, this scenario amounts to *c*_11_ > *c*_1*h*_ > *c*_12_, *c*_22_ > *c*_*h*2_ > *c*_12_ and *c*_*hh*_ > 0. For negative assortative mating, we have *c*_11_ < *c*_1*h*_ < *c*_12_, *c*_22_ < *c*_*h*2_ < *c*_12_, and *c*_*hh*_ < 0. The variance of X-chromosomal admixture in males depends on the contribution parameters and the limiting admixture in females, but not on the assortative mating parameters beyond their role in female admixture (eq. (47)). Therefore, we consider the variances only of autosomal admixture and X-chromosomal admixture in females.

#### 6.2.1 Positive assortative mating for each population

Here, we show that for positive assortative mating, eq. (45) can be written as the sum of a positive quantity and the limit of the variance of admixture for autosomes under random mating. That is,

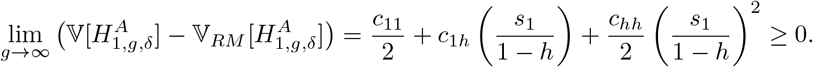

We recall that the quantity 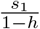 is the limit of the expectation of autosomal ancestry, and therefore takes its value in (0, 1) (VERDU and ROSENBERG, 2011, eq. 31). We can rewrite the right-hand side of eq. (48) as

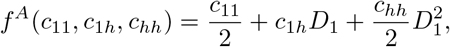

where *D*_1_ is a given constant in (0, 1). Under this assortative mating scenario, we can rewrite eq. (6) to give a range for *c*_1*h*_ of 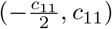 and, by our definition, we have *c*_11_, *c*_*hh*_ > 0 for positive assortment. Because *f* is linear in *c*_1*h*_, its minimum in terms of *c*_1*h*_ occurs at the boundary of the range of *c*_1*h*_. Substituting for *c*_1*h*_ its lower bound, 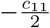, we have

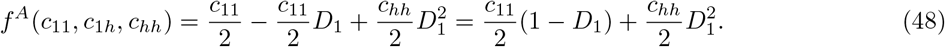

This quantity is positive because *D*_1_ takes its values in (0, 1), and *c*_11_, *c*_*hh*_ > 0. Therefore, we have demonstrated that this scenario of positive assortative mating increases the variance of autosomal admixture relative to a randomly mating population with the same contribution parameters.

Similarly, for X-chromosomal ancestry in females, we can show that for positive assortative mating, eq. (46) can be written as the sum of a positive quantity and the limit of the variance of the female X-chromosomal admixture under random mating. That is,

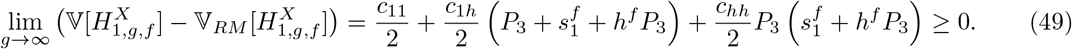

We recall that the quantities *P*_3_ and 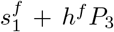 are the limits of the expectations of female X-chromosomal, and male X-chromosomal ancestry, respectively, and therefore take their values in (0, 1) (GOLDBERG and ROSENBERG, 2015, Appendix). We can rewrite eq. (49) as

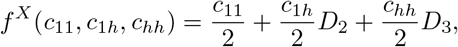

with *D*_2_ ∈ (0, 2) and *D*_3_ ∈ (0, 1). These upper bounds arise from the fact that 0 < *P*_3_ < 1 and 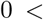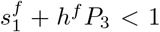 (GOLDBERG and ROSENBERG, 2015, Appendix). As in the cause for autosomal admixture above, *f* is linear in *c*_1*h*_. Therefore, its minimum in terms of *c*_1*h*_ occurs at the boundary of the range of *c*_1*h*_. Substituting for *c*_1*h*_ its lower bound, 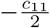, we have

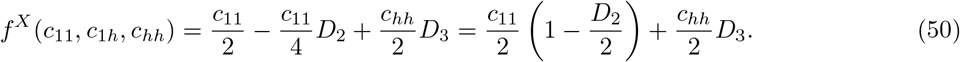

For this scenario of positive assortative mating, we can see that *f* is always positive because *c*_11_, *c*_*hh*_ > 0 by definition, and *D*_2_ ∈ (0, 2) and *D*_3_ ∈ (0, 1).

#### 6.2.2 Negative assortative mating for each population

A similar argument can be made for negative assortative mating. Under a scenario of intermediate preference for hybrids, we can rewrite eq. (6) to give a range for 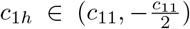 and *c*_*11*_, *c*_*hh*_ < 0 for negative assortment. To show that the maximum value *f*^*A*^ can take is less than zero, we use the upper bound of *c*_1*h*_, which is equal to 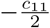, and is a positive value because *c*_11_ < 0. Using this bound gives the same quantity seen in eq. (48); under negative assortative mating, *f*^*A*^ is negative because *D* takes its values in (0, 1), and *c*_11_, *c*_*hh*_ < 0. Next, considering female X-chromosomal admixture, we can see that *f*^*X*^ is negative for all parameter values under this scenario of negative assortative mating. Substituting the lower bound of 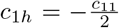 into (49), we have the same quantity seen in (50). For negative assortative mating, we have *c*_11_, *c*_*hh*_ < 0, and *D*_2_ ∈ (0, 2), *D*_3_ ∈ (0, 1). Therefore, *f*^*X*^ is negative for all parameter values in this negative assortative mating scenario.

In sum, when all populations experience positive assortative mating and the preference for one’s own population is the strongest preference, we see that the variances of autosomal and X-chromsomal ancestry can be written as a positive term plus the corresponding variance for a randomly mating population. Similarly, for negative assortment, the variances in equations (45) and (46) can be written as a negative term plus the variance in a corresponding randomly mating population. That is, holding the contributions 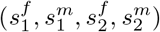 constant, positive assortative mating increases the variance of admixture and negative assortative mating decreases the variance relative to random mating.

#### 6.2.3 Positive mating with respect to *S*_1_

However, we can think of a more general type of assortative mating instead, allowing assortative mating to be positive in one population and random or negative in the other two populations. That is, we consider positive assortative mating with respect to population *i* as *c*_*ii*_ > *c*_*ij*_, *c*_*ji*_ for *i* ≠ *j*, and negative assortative mating with respect to population *i* as *c*_*ii*_ < *c*_*ij*_, *c*_*ji*_ for *i* ≠ *j*. Notably, these conditions are not mutually exclusive; positive assortative mating might occur in one population at the same time as negative assortative mating in another population. That is, in contrast to the assortative mating scheme above, we allow cases in which *c*_11_ > 0 while *c*_22_ < 0 within the definition of positive assortative mating *with respect to S*_1_.

Under this type of positive assortative mating, we allow cases in which *c*_11_ > *c*_12_ > *c*_1*h*_, and expand the range of *c*_1*h*_ to include 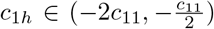 with *c*_11_ > 0. Therefore, eqs. (48) and (50) are no longer strictly positive, and they depend on the relative values of the assortative mating parameters as well as on the contribution parameters producing the expectation of admixture. Similarly, for negative assortative mating, 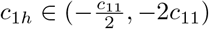 with *c*_11_, *c*_*hh*_ < 0, and eqs. (48) and (50) are not strictly negative.

Figure 6 plots an example of positive assortative mating for individuals from *S*_1_ and *H*, but negative assortative mating in *S*_2_. First, in green, we see an example with positive assortative mating in all populations, with *c*_11_ = *c*_*hh*_ = 0.02 and *c*_22_ = 0.06. In this scenario, the variance of admixture is higher than in the corresponding randomly mating population (black dashed line). Maintaining *c*_11_ = *c*_*hh*_ = 0.02, but now setting *c*_22_ = −0.04, the purple line plots a case in which positive assortative mating in *S*_1_ is overwhelmed by negative assortative mating in *S*_2_, producing a variance of admixture that is less than in the randomly mating population with the same contribution parameters.

**Figure 6:**
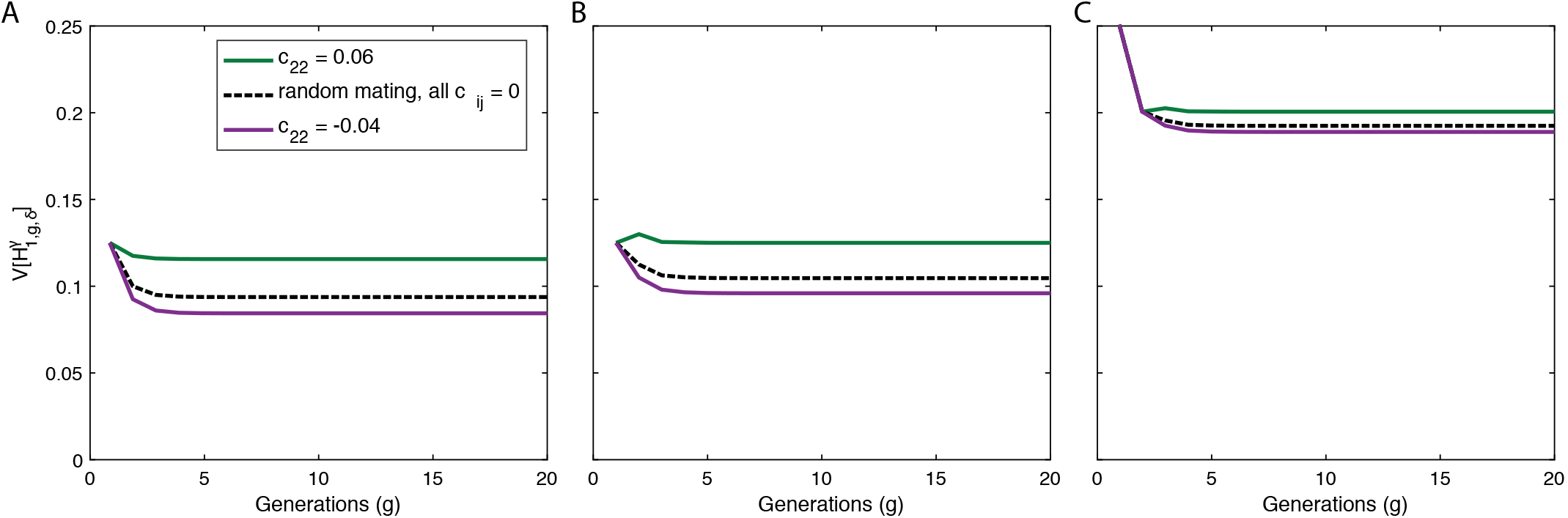
Example of the variance of ancestry under a constant-admixture scenario with assortative mating in producing *g* ≥ 2 when populations are allowed to differ in the direction of their mating preference. (A) Autosomes, 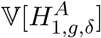. (B) X chromosomes in a female, 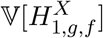. (C) X chromo-somes in a male, 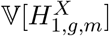. In all scenarios, populations *S*_1_ and *H* experience positive assortative mating, *c*_11_, *c*_*hh*_ = 0.02. Green curves show a scenario in which *S*_2_ also experiences positive assorta-tive mating, *c*_22_ = 0.06; purple curves show a scenario in which *S*_2_ experiences negative assortative mating, *c*_22_ = −0.04. The scenario of random mating with the same contribution parameters is the black dashed line. The plots use eqs. (20), (21), and (29)–(32). We set 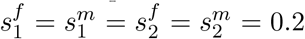, and use initial conditions 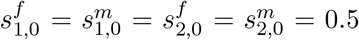 and *c*_11,0_ = 0. In this example, negative assortative mating in *S*_2_ offsets the positive assortative mating in *S*_1_ and *H*, producing lower variances of admixture in *S*_1_, 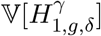, in the assortatively mating population than in the randomly mating population.

## 7 Discussion

### 7.1 Summary of results

A recent series of papers has devised mechanistic models to consider admixture over time (VERDU and ROSENBERG, 2011; GOLDBERG *et al*., 2014; GOLDBERG and ROSENBERG, 2015), assuming that the parents of a randomly chosen individual from the admixed population are chosen independently. We have extended these models to consider the effect on the distribution of admixture of preferential mating based on population of origin. Although the mean autosomal and X-chromosomal admixture fractions depend only on the sex-specific contributions from the source populations (eqs. (11) and (25)), we have found that the variances of admixture contain signatures of assortative mating (eqs. (21), (31), and (32)).

For both autosomes and X chromosomes, when all three populations experience positive assortative mating, the variance of admixture always increases relative to a randomly mating population with the same contributions (Fig. 5). Similarly, when all three populations experience negative assortative mating, the variance of admixture always decreases. Because an individual’s autosomal admixture fraction is the average of its parents’ admixture fractions, in the absence of new contributions from the source populations, random pairings of different admixture fractions decrease the variance of admixture every generation (GOLDBERG and ROSENBERG, 2015). However, positive assortative mating decreases the proportion of matings that occur between different admixture fractions, with more low-low and high-high pairings maintaining a wide spread in the distribution of admixture. Similarly, negative assortative mating increases the proportion of matings that occur between pairs with different admixture fractions, thereby decreasing the spread of the admixture fraction distribution.

Assortative mating has a smaller effect on the variance of X-chromosomal admixture than on the variance of autosomal admixture. Conditional on the distribution of ancestry in the previous generation, the variance of X-chromosomal admixture in a male chosen at random from the admixed population is equivalent for randomly mating and assortatively mating populations (eq. (40)) because it is determined by the female X-chromosomal ancestry only, not by the ancestries of both members of a mating pair. Hence, the effect of assortative mating on the variance of X-chromosomal admixture is seen only in females (eq. (39)).

Our model is flexible, allowing assortative mating to occur in different directions and strengths in the three populations. That is, positive assortative mating with respect to *S*_1_ can occur at the same time as random mating or negative assortative mating in *H* and *S*_2_. We found that the effect of assortative mating on the variance of admixture depends on trade-offs in the preferences among the three populations. For both autosomes and the X chromosome, certain scenarios of assortative mating with different directions of mating preference in different populations change the direction of the effect of assortative mating on the variance of admixture (Fig. 6).

### 7.2 Comparisons to previous models of assortative mating

Focusing on admixed populations, we chose to model mating preferences by population of origin. This type of assortative mating captures scenarios in which the admixed population is a distinct group, and preferences follow population membership rather than any specific trait, ancestry, or genotype. Different populations might have different geographical locations, trait preferences, or host preferences. For example, scenarios of preference for or against new migrants compared to other admixed individuals have been hypothesized in swordfish, butterflies, and grasshoppers (RITCHIE *et al*., 1989; HOWARD, 1993; DUENEZ-GUZMAN *et al*., 2009; MELO *et al*., 2009; SCHUMER *et al*., 2017). Similarly, in admixed human populations, language barriers or social identities may produce population-based preferences (QIAN, 1997; JACOBSON *et al*., 2004; RUIZ-LINARES *et al*., 2014).

Our model of assortative mating by population recapitulates patterns seen in other assortative mating models in settings that do not consider admixture. Increased variance in a phenotype, with no change in the mean, has been observed in classic models of positive assortative mating by a single-locus genotype or a quantitative trait value without admixture (JENNINGS 1916, pgs. 66-68; FISHER 1918, pgs. 410-414; WRIGHT 1921, pgs. 153-155; CROW and KIMURA 1970, ch. 4). These studies demonstrate that under a model of assortative mating by genotype, positive assortative mating produces an increase in the proportion of homozygotes compared to heterozygotes, analogous to the excess proportion of admixture fractions equaling zero and one seen in our model. For the X chromosome, studies of the distribution of traits and the correlation of relatives under assortative mating by trait loci also find that assortative mating influences X chromosomes in females more than in males (RISCH, 1979; YENGO and VISSCHER, 2018).

Empirical examples of the correlation in ancestry between members of mating pairs in admixed populations motivated ZAITLEN *et al*. (2017) to model mating preferences as a fixed correlation in admixture fraction between mates in the admixed population. Extending a Wright-Fisher model, ZAITLEN *et al*. (2017) found a similar inflation of the variance of admixture under positive assortative mating and a decrease in the variance of admixture with negative assortment. In their model, mating preferences have the same direction and strength for all individuals; therefore, their model does not allow for the cases we examine in which the variance of admixture decreases under positive assortative mating with respect to some populations and negative assortative mating in others.

### 7.3 Sex-biased admixture and assortative mating

Here, we assumed equal female and male contributions from each of the two source populations, 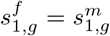 and 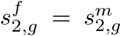 for some of our analyses. However, sex bias and assortative mating may occur in the same population (BRYC *et al*., 2015; ZOU *et al*., 2015). Our work does allow consideration of different sex-specific contributions from the sources. GOLDBERG and ROSENBERG (2015) examined the effect of sex-biased admixture on the mean X-chromosomal and autosomal admixture in the case of random mating, *c*_ij_ = 0 for all *i* and *j*. When estimating sex bias levels from data, because assortative mating does not influence the mean admixture levels, inclusion of assortative mating would not change estimates of sex bias obtained from mean X-chromosomal and autosomal admixture levels.

### 7.4 Assortative mating can bias inference of the timing of admixture

The variance of admixture is informative about the timing of admixture (VERDU and ROSENBERG, 2011; GOLDBERG *et al*., 2014; LIANG and NIELSEN, 2014). Because assortative mating often changes the variance of admixture, estimating the timing of admixture without accounting for assortative mating may lead to bias in estimates of the timing of admixture.

For example, considering autosomal admixture under the case of a single admixture event, we can rewrite eq. (41) to consider how assortative mating affects this inference in the case of a single admixture event. For an observed variance of autosomal admixture *V*^A^, we have

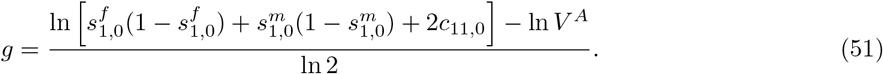

The estimated timing of admixture, *g*, is directly related to the assortative mating constant *c*_11,0_. Therefore, fixing 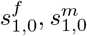, and *V*^*A*^, positive assortative mating increases *g* relative to random mating. That is, failing to account for positive assortative mating makes admixture appear more recent than the true admixture event. This observation follows from the increase in variance under positive assortative mating compared to random mating: because the variance decreases over time, a fixed variance persisting due to positive assortative mating might be misinterpreted as a more recent admixture under random mating. Negative assortative mating has the opposite effect, in that failing to account for negative assortative mating produces an overestimate of *g* relative to the true level.

Figure 7 plots estimates of *g* according to eq. (51) for varying levels of *c*_11,0_, considering both positive and negative assortative mating, specifying *V*^*A*^ = 0.0625, and 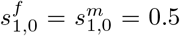 under a single admixture event. For a randomly mating population, an autosomal variance of 0.0625 implies an admixture event 2 generations ago under a single-admixture model with 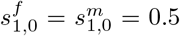. However, when assortative mating is allowed, the admixture event might have taken place in the current generation or up to 3 generations ago, with the estimated *g* monotonically increasing in *c*_11,0_. When *c*_11,0_ ≥ 0, the age of admixture is greater than in a randomly mating population with the same variance and contributions. Conversely, when *c*_11,0_ ≤ 0, estimates of *g* are smaller than in a corresponding randomly mating population.

**Figure 7:**
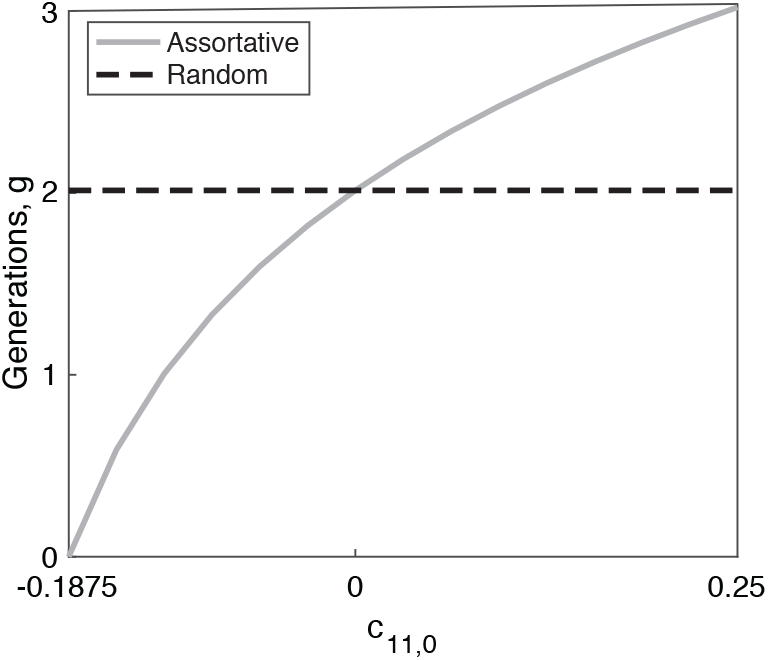
Timing of admixture under assortative mating. The number of generations since admixture, *g*, for a single admixture event is plotted as a function of *c*_11,0_ using the variance in autosomal admixture, eq. (51), with *V*^*A*^ = 0.0625 and 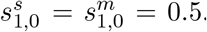. The plot traverses the range of possible *c*_11,0_ values for *g* ≥ 0. Recalling that the values of the *c*_*ij*,0_ are bounded such that the probability of each given parental pairing takes its values in the interval [0, 1], and such that each probability is no greater than the probability of one of its constituent components, for *c*_11,0_ > 0, we have 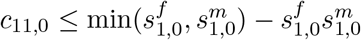. In this case, *c*_11,0_ ≤ 0.25. For *c*_11,0_ < 0, we have 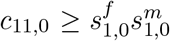. In this case, we have *c*_11,0_ ≥ −0.25. However, *g* cannot be negative; therefore, in this case, we truncate the domain at the value of *c*_11,0_ that produces *g* = 0. The value of *g* for a randomly mating population with the same contributions and variance is shown in the dashed line.

The variance-inflating effect of positive assortative mating on the admixture level is similar to that described by ZAITLEN *et al*. (2017) for the magnitude of ancestry linkage disequilibrium. However, because different scenarios of assortative mating affect the variance of admixture in different directions, under our model, we show that failing to account for assortative mating can either underestimate or overestimate the timing of admixture, depending on whether assortative mating is positive or negative.

## Acknowledgments

We acknowledge support from NIH R01 HG005855 and NSF BCS-1515127 to NAR.

## Appendices

## A Extrema of 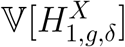 for a single admixture event in a randomly mating population

This appendix derives the extrema of 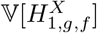 and 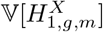 as functions of the contribution parameters 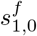 and 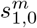 for a randomly mating population founded in a single admixture event with no further contributions. The corresponding extrema for the variance of autosomal admixture were previously analyzed by GOLDBERG *et al*. (2014). Here, we use the model of GOLDBERG and ROSENBERG (2015) to study the variance of X-chromosomal admixture in a randomly mating population (*c*_11,0_ = 0). We perform this computation for completeness, to provide parallel results for the X-chromosomal case to those derived by GOLDBERG *et al*. (2014) for the autosomes.

Figure A1 plots the variance of X-chromosomal admixture in females using eq. (44) with *c*_11,0_ = 0, for multiple values of *g*. As *g* increases, the admixed population mixes with no further contributions from the source populations, and the variance decreases for all values of 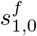 and 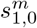.

From eq. (44), at *c*_11,0_ = 0, the maximum of 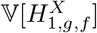 in terms for 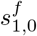 and 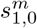 occurs when 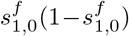 and 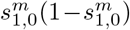 are separately maximized. That is, the maximum occurs when 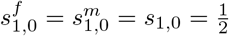.

For the minimal variance, which equals 0, equation (44) equals 0 at four points: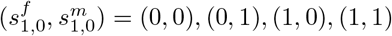. The locations of the global maximum and the four minima are the same as those seen for the autosomal variance by GOLDBERG *et al*. (2014). If only a single population contributes to the hybrid population 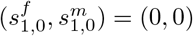, or (1, 1), then no variation exists in the population because all individuals have the same admixture fraction of 0 or 1. Similarly, when all males are from one source population and all females are from the other, so that 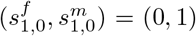 or (1, 0), no variation exists because all individuals have an admixture fraction of 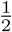.

Next, we can find the critical points of the X-chromosomal variance for a given value of *s*_1,0_, permitting 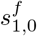 and 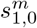 to vary. Recalling eq. (2), we rewrite the variance of X-chromosomal admixture (eq. (44), with *c*_11,0_ = 0) as a function of 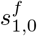 and *s*_1,0_. We have

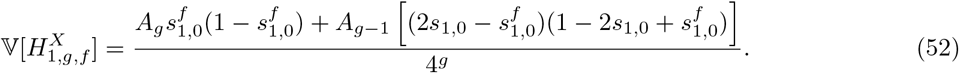

Setting the first derivative of eq. (52) with respect to 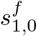 equal to 0, we find that the critical point of the variance of X-chromosomal admixture occurs at

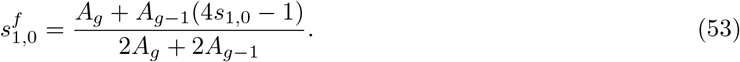

This critical point is a maximum because the second derivative of eq. (52) is negative, −2(*A*_*g*_ + *A*_*g*−1_).

Following the same procedure, we can write the variance of X-chromosomal admixture in females as a function of 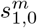 and *s*_1,0_. We have

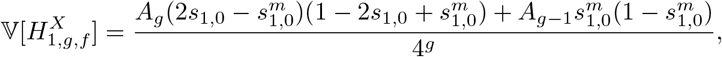

producing a maximum at

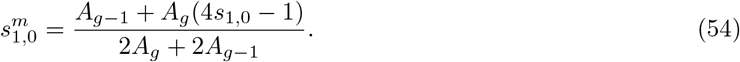

**Figure A1:**
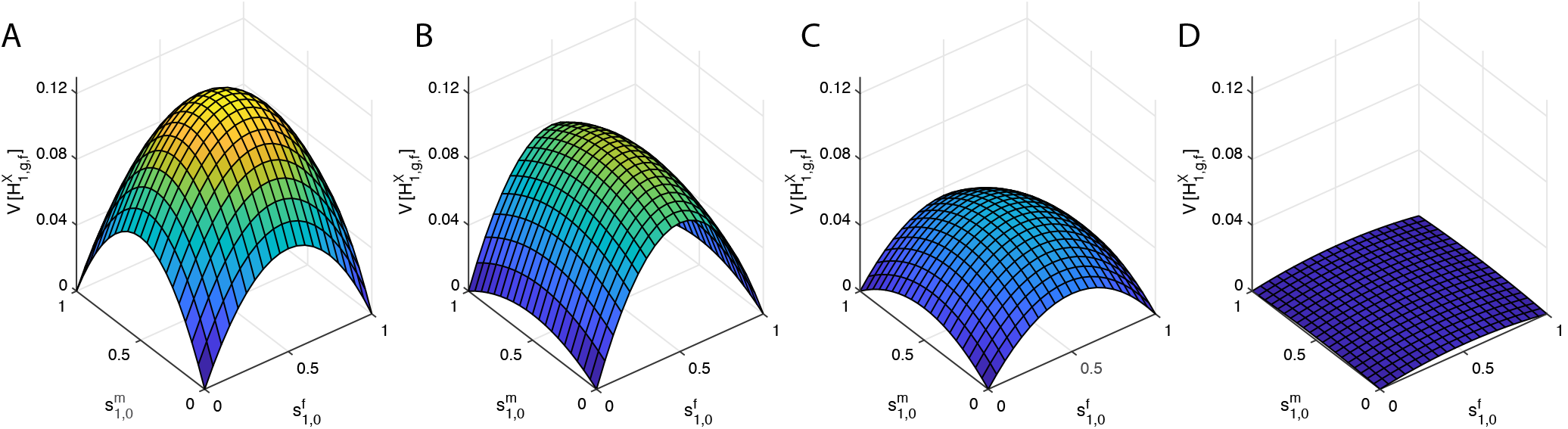
The variance of ancestry as a function of the sex-specific contributions for a single admixture event in a randomly mating population. (A) *g* = 1. (B) *g* = 2. (C) *g* = 3. (D) *g* = 8. In each panel, 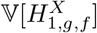 is plotted over the range of permissible values of 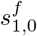 and 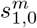, using eq. (44).

We also have the constraints that 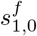 and 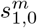 take their values in [0, 1]. eqs. (53) and (54) lie within this interval when both of the following pairs of inequalities hold:

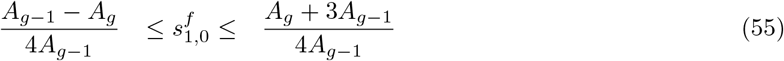

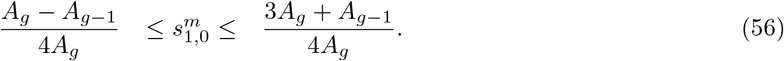

Eq. (55) is always satisfied, as *A*_*g*_ > *A*_*g*−1_, so that the left hand side of the inequality is negative and the right hand side exceeds 1. Therefore, the maximum always occurs when 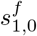 takes the value in eq. (53). When eq. (56) is satisfied, the maximum occurs when 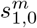 follows eq. (54), and when 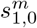 is outside the bounds in eq. (56), the maximum occurs at 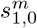 equal to 0 or 1.

Next, we find the minima of the variance of the female X-chromosomal admixture fraction with respect to 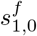 and 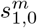 for specified *s*_1,0_. Eqs. (53) and (54) are quadratic with a single critical point that is a maximum. Therefore, the minima of the variance of X-chromosomal admixture must lie along the boundary of the interval in which 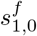 and 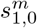 take their values, [0, 1].

For *s*_1,0_ ∉ {0, 1}, the minimum variance is on the boundary of parameter values, 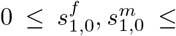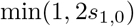. Specifically, for 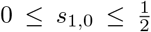, the minimum variance occurs when 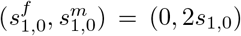. Using eq. (44), we see that because *A*_*g*_ > *A*_*g*−1_, the variance of the female X-chromosomal admixture fraction is smaller when 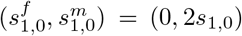 than when 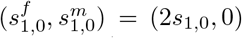. Therefore, unlike for the autosomes, the minima are not symmetric for the X chromosome (Figure A1, for *g* > 1). Specifically, because *A_g_* is monotonically increasing, variance depends to a greater extent on 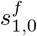 than on 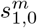 and the minimum variance for a specific *s*_1,0_ occurs when *s*_1,0_ = 0, but not 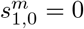. Similarly, for 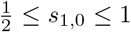, the minimum variance occurs when 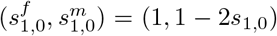. Note that for 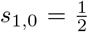 both 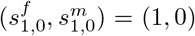 and 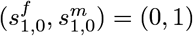 are minima.

## B Closed-form expression and temporal limit for 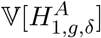 under constant contributions over time in an assortatively mating population

In this appendix, we use the closed form of the variance of autosomal admixture for a randomly mating population, derived by GOLDBERG *et al*. (2014), to write an expression for 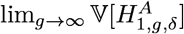 under assortative mating.

GOLDBERG *et al*. (2014) derived expressions for 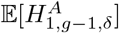 and 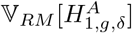 (eq. 37 and eqs. 51–52, respectively). We substitute these equations in eqs. (22) and (23) to produce a closed form for the variance under assortative mating by population. We have

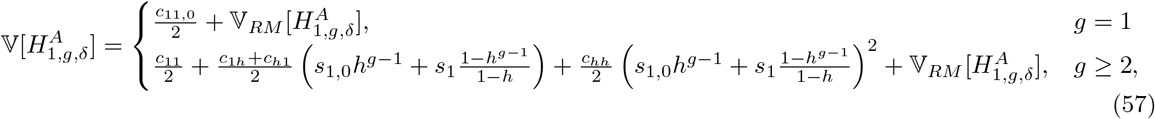

where 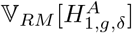 is given by eqs. 51 and 52 from GOLDBERG *et al*. (2014).

For the limit as *g* → ∞, we recall eq. 53 from GOLDBERG *et al*. (2014),

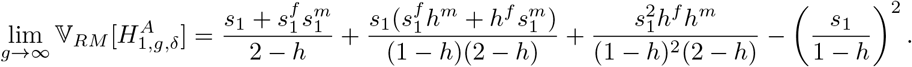

Therefore, taking the limit as *g* → ∞ of the variance of autosomal admixture in eq. (57), we have

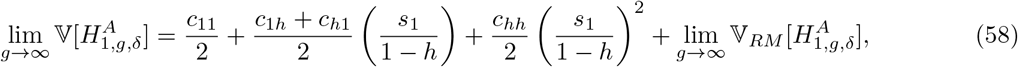

Finally, assuming assortative mating is the same for the two sexes, *c*_*ij*_ = *c*_*ji*_, we have eq. (45).

## C Closed form expression and temporal limit for 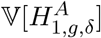 under constant contributions over time in a randomly mating population

Here, we find an expression for the limit of the variance of X-chromosomal admixture as *g* → ∞ in a randomly mating population. We follow the structure of Appendix 1 from GOLDBERG and ROSENBERG (2015). Using eqs. 3–4 from GOLDBERG and ROSENBERG (2015) and eqs. (33)–(35) above, we can rewrite the variance of the X-chromosomal admixture fraction in females for the case of constant admixture. For *g* = 1, we have equation (33). For *g* = 2, we have

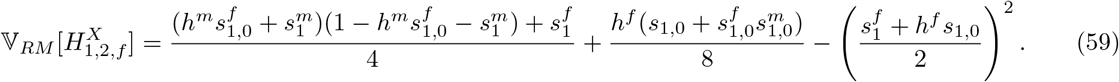

For *g* ≥ 3, we rewrite the female X-chromosomal variance as a second order recursion using equations (35) and (36). We have

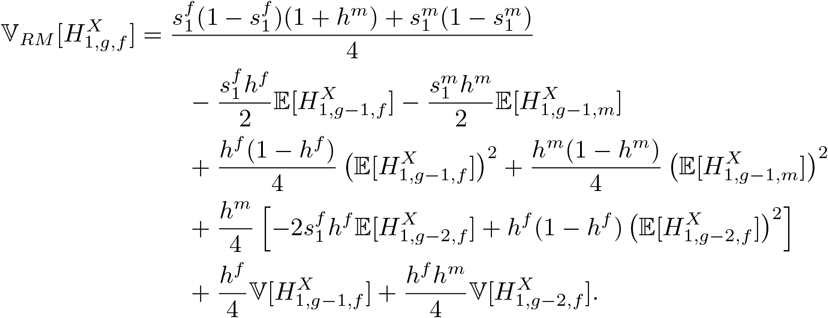

We simplify the notation by defining 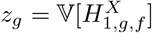. For *g* ≥ 3, we have

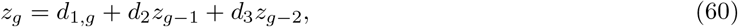

with

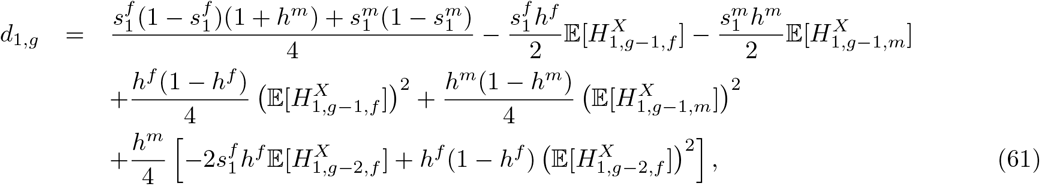

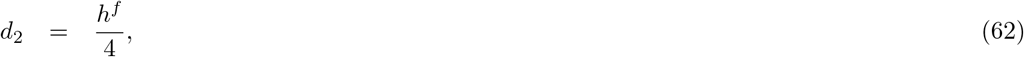

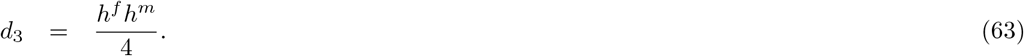

In eq. (61), *d*_1,*g*_ depends on *g* through the expectation of X-chromosomal admixture. However, we can use closed-form solutions for 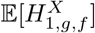 and 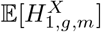 (GOLDBERG and ROSENBERG, 2015, eqs. 17 and 18).

From eqs. (33) and (59), respectively, we have

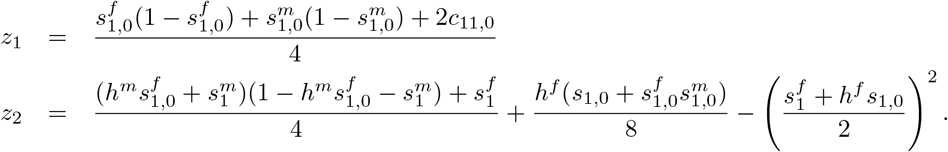

We define a generating function 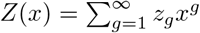 whose coefficients *z*_*g*_ represent the values of 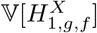 in each generation. As the admixed population does not yet exist in generation 0,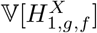 and *Z*(*x*) are undefined for *g* = 0. For convenience, we therefore work with *W* (*x*) = *Z*(*x*)*/x*, setting *w*_*g*_ = *z*_*g*+1_ for *g* ≥ 0. We then have

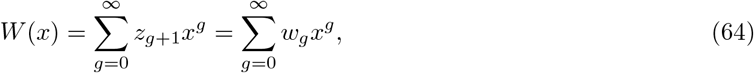

and *w*_*g*_ = *d*_1,*g*_ + *d*_2_*w*_*g*−1_ + *d*_3_*w*_*g*−2_ for *g* ≥ 2 by eq. (60). Using eq. (64), it follows that

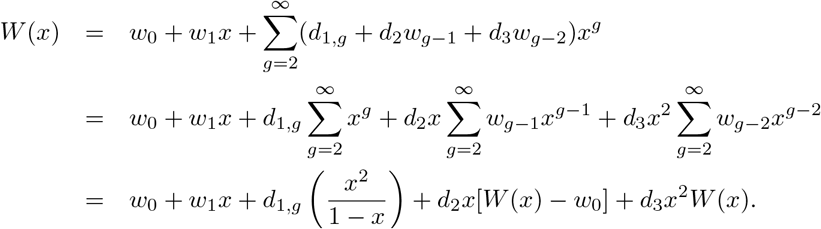

Solving for *W* (*x*), we have

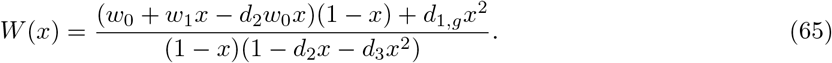

We can decompose the expression in eq. (65), producing

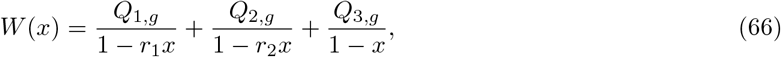

where *r*_1_ and *r*_2_ are reciprocals of the two roots of 1 − *d*_2_*x* − *d*_3_*x*^2^,

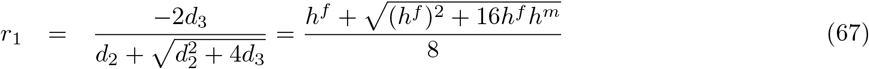

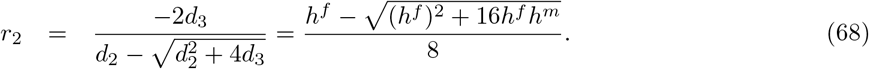

Setting eq. (65) equal to eq. (66), we have

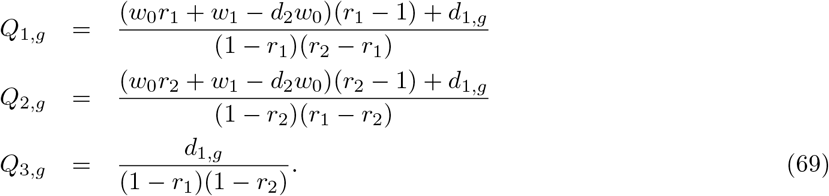

The Taylor expansion of eq. (66) around *x* = 0 then gives

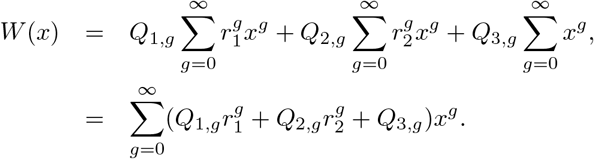

Therefore, for *g* ≥ 0,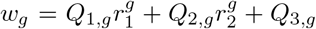, and the closed-form expression for the X-chromosomal female mean admixture fraction in generation *g* ≥ 1, 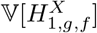, is

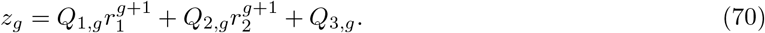

Because for *h*^*f*^, *h*^*m*^ ∈ [0, 1], *r*_1_ monotonically increases in *h*^*f*^ and *h*^*m*^ and *r*_2_ monotonically decreases, the maxima and minima of *r*_1_ and *r*_2_ over permissible values of *h*^*f*^ and *h*^*m*^ occur at the boundaries of the closed interval [0, 1]. Using eqs. (67) and (68), we have 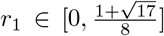 and 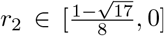; we exclude 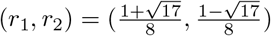 as *h*^*f*^ and *h*^*m*^ cannot both be 1. Because |*r*_1_|, |*r*_2_| < 1, the variance of the female X-chromosomal admixture fraction in eq. (70) approaches a limit as *g* → ∞. Using eqs. (70) and (36), we can find expressions for the limit of the variance of the X-chromosomal admixture fractions,

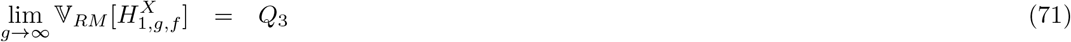

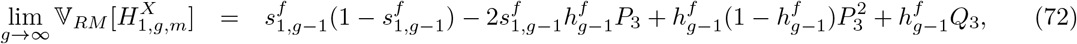

where *P*_3_ is the limit of the mean female X-chromosomal admixture fraction (GOLDBERG and ROSENBERG, 2015, eq. 19), and *Q*_3_ = lim_g→∞_ Q_3,g_. Rewriting *P*_3_ here, we have

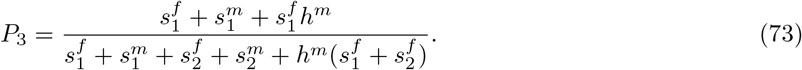

*Q*_3_ is obtained by substituting 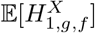 and 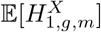 with their limits, *P*_3_ and 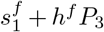, respectively, into eq. (69). Using eqs. (61)–(63) to simplify, we have

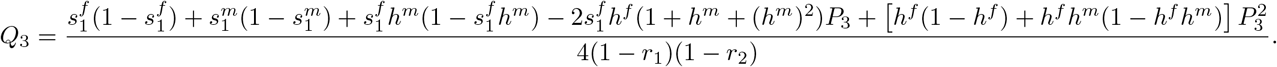

